# TRiC activates the unfolded protein response and protects starved stem cells by modulating energy and lipid metabolism during planarian regeneration

**DOI:** 10.1101/732875

**Authors:** Óscar Gutiérrez-Gutiérrez, Daniel A. Felix, Alessandra Salvetti, Anne Thems, Stefan Pietsch, Andreas Koeberle, K. Lenhard Rudolph, Cristina González-Estévez

## Abstract

Fasting protects stem cells and increases stem cell functionality through mechanisms which are not fully understood. Planarians are not only able to regenerate their bodies, but also to stand long periods of starvation by shrinking in size. This adaptation is possible because of a large population of adult stem cells which indefinitely self-renew even under starved conditions and thus confer planarians with immortality. How starved planarians are able to maintain healthy stem cells and to fuel stem cell proliferation allowing regeneration is unknown. Here we found the TCP-1 ring complex (TRiC) to be upregulated in starved stem cells. Down-regulation of TRiC impairs planarian regenerative response by inducing stem cell genome instability, mitotic defects and stem cell death which translates into stem cell exhaustion. This regulation is specific of starvation since feeding planarians prevents the phenotype. Importantly we found that TRiC activates the unfolded protein response (UPR) which allows a convergent regulation of cellular energy and lipid metabolism in starved planarians thus permitting the high energy demanding regenerative mitotic response. We identified a novel mechanism through which starvation protects the somatic stem cell genome allowing for unlimited stem cell proliferation and regeneration.

## Introduction

Fasting increases lifespan and healthspan in many organisms (Fontana and Partridge, 2015; Longo and Mattson, 2014). This is due at least in part to an increase in stem cell functionality which decreases during ageing and contributes to age-associated pathologies and the overall ageing process (Adams et al., 2015; Behrens et al., 2014). Indeed diet is emerging as an important regulator of adult stem cell function. For instance, dietary restriction (DR) improves the repopulation capacity of hematopoietic stem cells (HSCs) in early mouse ageing (Tang et al., 2016) and enhances stem cell functionality in intestinal epithelium (Yilmaz et al., 2012) and muscle (Cerletti et al., 2012). It is also suggested that fasting confers stem cell protection, since cycles of prolonged fasting and refeeding prior to chemotherapy protects hematopoietic stem cells (HSCs) but not cancer cells from DNA damage thus promoting their self-renewal and regeneration (Bauersfeld et al., 2018; Lee et al., 2012; Nencioni et al., 2018; Raffaghello et al., 2008). Our understanding on the mechanisms that contribute to this protective and pro-regenerative effect and mediate the beneficial effects of DR and fasting on stem cell function is limited to few signalling pathways (reviewed in (Mihaylova et al., 2014).

Proteome integrity is essential for the correct functionality of cells and the survival of organisms. The maintenance of protein homeostasis (proteostasis) is regulated by a complex network, which coordinates protein synthesis, folding, trafficking, aggregation, disaggregation and degradation of proteins. This balanced network is constantly affected by environmental factors such as mutations and age. Indeed the ability of post-mitotic cells to maintain proteostasis is gradually compromised with age and this associates with many human age-related pathologies, such as neurodegenerative disorders, cardiovascular diseases and cancer (Balch et al., 2008; Hipp et al., 2019; Lindquist and Kelly, 2011; Morimoto, 2008). Interestingly, interventions that increase lifespan such as DR or reduction of insulin/insulin-like growth factor 1 (IGF-1) signalling are associated with enhanced mechanisms regulating proteostasis in postmitotic cells, in most cases through a reduction in protein synthesis and the activation of protein degradation by autophagy and the proteasome (Cohen et al., 2009; Vilchez et al., 2012b).

Chaperones are another component of the proteostasis network and are responsible for assisting in *de novo* folding and for protecting of existing proteins from proteotoxic stress (Labbadia and Morimoto, 2015). There are specific chaperones for each of the protein-folding compartments in a cell (e.g. the cytosol, the endoplasmic reticulum (ER) and the mitochondria) that respond to compartment-specific pathways, the cytosolic stress response and the unfolded protein response pathways of the ER (UPR^ER^) and mitochondria (UPR^mt^) (Buchberger et al., 2010) to overcome different stressors. The cytosolic stress response induces the expression of chaperones, cochaperones and chaperonins under different sorts of stress that are categorized as ATP-dependent (chaperonins/HSP60s, HSP70s, HSP90s and HSP100s) or ATP-independent (small HSPs). Chaperones act alone or in combination with various cochaperones to regulate client-substrate interactions, folding, disaggregation, degradation and trafficking within the cell. Compared to other chaperones, the HSP60/chaperonin member TRiC (TCP1-ring complex or chaperonin containing TCP1, also known as CCT) recognizes a smaller repertoire of substrates and is necessary for folding about 5-10% of newly synthesized proteins including actin and tubulin (Saibil, 2013). It also binds to misfolded proteins regulating their aggregation. Indeed it has been predicted that late folding intermediates or misfolded species are preferred substrate conformers of TRiC (Horwich et al., 2007). The ER is the major organelle for lipid synthesis and the biosynthesis/folding and maturation of proteins. When overburdened by misfolded proteins or exceeded in the capacity to export proteins and lipids, the UPR is activated at the ER, which decreases global protein translation and induces the specific transcription of genes to alleviate stress and restore proteostasis to promote cell survival. Misfolded, aggregated or damaged proteins are degraded through the proteasome or autophagy. If the UPR fails to restore the ER to normality, ER stress can promote apoptosis (Labbadia and Morimoto, 2015).

Although all cells are able to activate these stress response pathways, it has been hypothesized that stem cells employ them more in order to protect their proteome and maintain their pluripotency and immortality (Noormohammadi et al., 2018; Vilchez et al., 2014). For instance, human embryonic stem cells (hESCs) present increased levels of some chaperones (Noormohammadi et al., 2016) and a higher proteasome activity (Vilchez et al., 2012a) compared to their differentiated counterparts that decreases during differentiation. Similarly, activated neural stem cells (aNSCs) have increased expression of proteasome-associated genes and some chaperones (Leeman et al., 2018).

While these studies show the importance of proteostasis regulation in stem cells, it remains unclear whether loss of proteostasis plays an important role in the decline of stem cell (i.e. active somatic stem cells) function observed during ageing and whether fasting- or DR-induced beneficial effects on stem cells involve a superior maintenance of proteostasis. Here we use the planarian species *Schmidtea mediterranea*, a consolidated model for the study of stem cells and regeneration (Reddien, 2018; Rink, 2013). Planarians possess a large population of somatic stem cells in their bodies which includes pluripotent stem cells (Wagner et al., 2011) and permits planarians to fully regenerate their bodies in few days. Remarkably, planarians are also able to stand starvation by shrinking in size (or “degrow”) while maintaining the relative number of stem cells and the same regenerative power as fed or growing planarians (Felix et al., 2019; Gonzalez-Estevez et al., 2012a). Since food availability fluctuates in nature, growth and degrowth are normal cycles in the planarian life (Felix et al., 2019; Pellettieri, 2019). Stem cell maintenance during starvation could be a strategy to allow for a rapid growth when a more favorable nutritional environment is encountered, or for being primed for a regenerative response to injury (Felix et al., 2019). Planarians are considered immortal, since their stem cell pool is able to infinitely replace old or damaged cells and to respond to injury (Felix et al., 2019; Sahu et al., 2017). Recently we have shown that starvation positively influences planarian stem cells, since it rejuvenates the stem cell pool in terms of telomere length through downregulation of mTOR signalling (Iglesias et al., 2019). However it is currently unknown how planarian stem cells under conditions of starvation are able to cope with the massive demand of proliferation required for regeneration (Baguñà, 1976b; Wenemoser and Reddien, 2010).

Here we find that mechanisms regulating proteostasis (i.e. TRiC and UPR) are upregulated in stem cells during starvation. Down-regulation of TRiC impairs the planarian regeneration response specifically in starved planarians since feeding prevents the TRiC RNAi phenotype. We show that TRiC is necessary to maintain stem cell genome integrity, allows mitosis and overall prevents stem cell exhaustion, an important driver of organismal ageing. We also find that TRiC activates the UPR which allows a convergent regulation of lipid metabolism and overall levels of cellular energy, which is specifically necessary for starved stem cells to be able to mount the regenerative response. We propose a model where stem cell-specific chaperones, through the UPR, protect stem cells during starvation and allow regeneration by mobilizing lipid droplets to provide energy and/or lipid membrane components necessary for the cell to grow and proliferate. Our findings suggest a novel cross-talk between the stress response at the cytosol and at the ER which would be necessary to maintain stem cell genome integrity and allow stem cell proliferation and regeneration in the immortal planarian during starvation. We also shed light into different regenerative programmes depending on the nutritional status. Our data contributes to the understanding of proteostasis in stem cell biology during fasting and ageing and highlights the importance of doing ageing research in long-lived animals.

## Results

### Transcriptional profiling reveals enhanced TRiC expression in stem cells in response to starvation

In order to investigate the transcriptional profiles of stem cells at different nutritional status we sorted stem cells by FACS (X1 subfraction; S and G2/M cell cycle phase stem cells) (Figure S1A) (Hayashi et al., 2006) from 1, 7 and 30 days post-fed or starved planarians (1dS, 7dS and 30dS, respectively) and performed RNA-seq and pairwise comparison between the different nutrient conditions. By performing gene ontology (GO) enrichment we found that “mitotic S phase” was among the most overrepresented biological processes down-regulated in X1 stem cells at 7dS and 30dS versus 1dS (Figure 1A-B and Table S1A-C). This suggests that the planarian X1 subfraction of stem cells is enriched in S phase in 1dS planarians when compared to planarians at 7dS and 30dS. This agrees with 1dS planarians being under the feeding mitotic peak (Baguñà, 1976) and with previous observations in other organisms under a feeding response (Duan et al., 2017) and thus supports that 1dS is a “feeding response” time point in planarians. The GO-term “gene expression” was also found among the down-regulated processes at 7dS and 30dS when compared to 1dS, suggesting that cycling stem cells in starved planarians down-regulate transcription and translation as described for many organisms under nutrient limitation (Hansen et al., 2007; Wullschleger et al., 2006) (Figure 1A-B and Table S1B-C). Interestingly we also found that stem cells at 7dS and 30dS up-regulate components of known pathways that promote increased stem cell functionality during DR in mammalian stem cells, such as the insulin signalling pathway (Mihaylova et al., 2014) (Table S1D).

**Figure 1.**
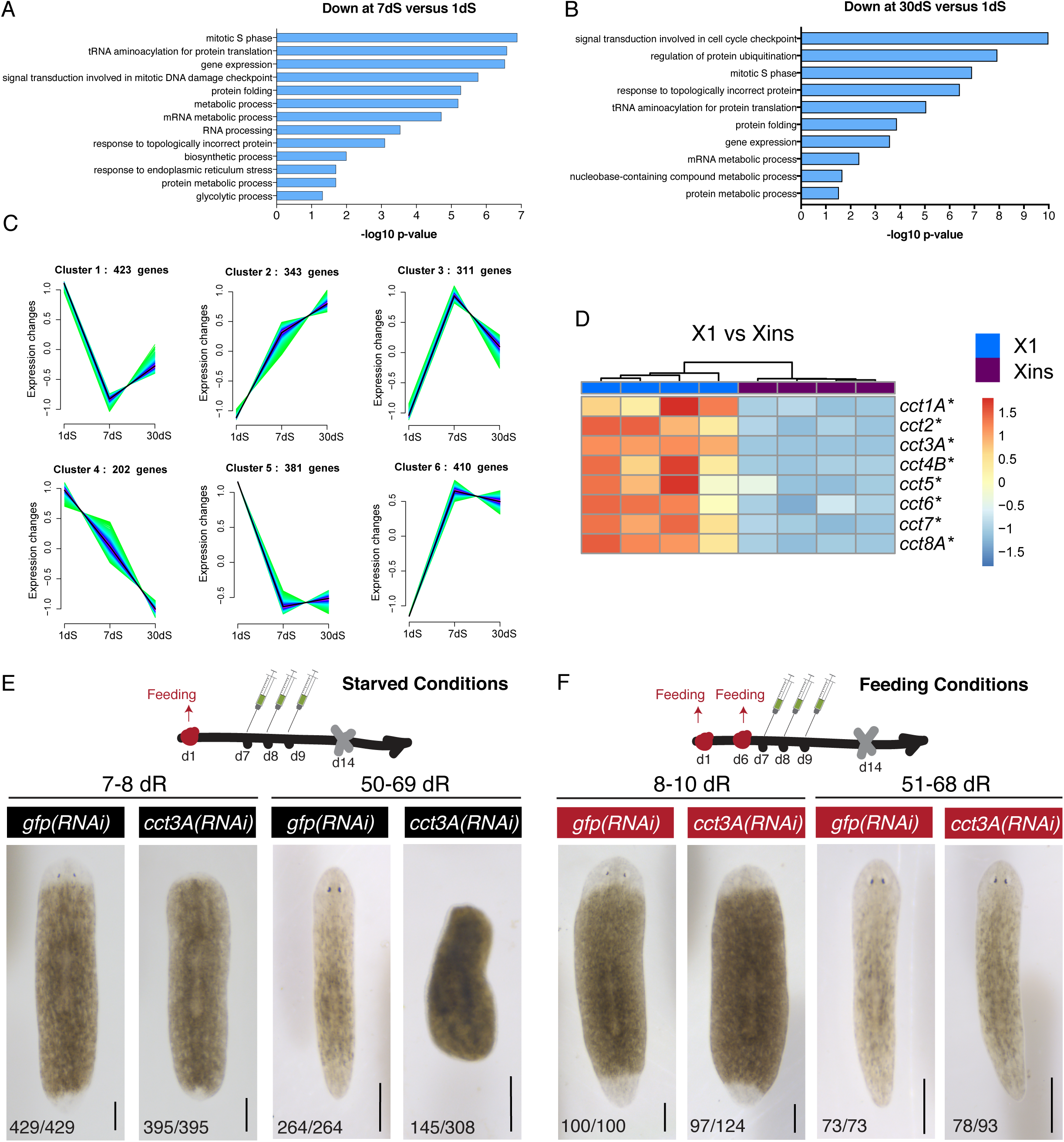
Transcriptional profile of stem cells (X1) at different nutritional status finds TRiC enriched in stem cells at 1dS and 30dS and *cct3A* necessary for blastema formation during starvation. (A) GO enrichment analysis for biological processes of the 814 down-regulated genes (q-value < 0.1) at 7dS versus 1dS in X1 (stem cells). (B) GO enrichment analysis for biological processes of the 491 down-regulated genes (q-value < 0.1) at 30dS versus 1dS in X1. No enriched biological processes were found up-regulated. For better visualization in A and B, similar enriched GO terms based on the same subset of genes were manually removed to reduce redundancy. (C) Clustering generated from 2070 DEGs found in stem cells (X1) according to their expression during different nutritional conditions (1dS, 7dS and 30dS). DEGs are grouped in 6 clusters with similar expression profiles. Cluster 1 is characterized by a U shape. Number of genes assigned to every cluster is showed on the upper part of every graph. (D) The expression heatmap shows that all the *cct* subunits from TRiC are up-regulated in X1 (stem cells) compared to Xins (differentiated cells) at 30dS. Colour-coded scale covers the range of TPM (transcripts per million) values for every replicate. (E) RNAi injections schedule in starved conditions. Planarians are at 14dS when the amputation is performed (indicated by the grey cross at day 14). Live images show that *cct3A(RNAi)* planarians form a minimal blastema compared to controls at the time points shown. At the bottom are the number of planarians with the phenotype shown. The remaining planarians are dead by the time point of regeneration shown. (F) RNAi injections schedule in feeding conditions. One extra feeding respect to E is introduced one day prior to injections. Planarians are at 8dS when the amputation is performed (indicated by the grey cross at day 14). The live images show that most of the *cct3A(RNAi)* planarians regenerate as controls. At the bottom are the number of planarians with the phenotype shown. The remaining planarians are either dead (4/124 at 8-10dR and 10/93 at 51-68dR) or have a tiny blastema and died later (23/124 at 8-10dR and 5/93 at 51-68dR). Scales, 300 µm.

Another overrepresented biological process that stirred our interest was “protein folding” (Figure 1A-B) with many of the transcripts corresponding to cytosolic chaperones up-regulated during the feeding response (1dS) when compared to 7d (25/41) and 30dS (12/21) (Table S1E-F). In addition to their functions in folding of *de novo* synthesized proteins, many chaperones are also induced under conditions of environmental stress and are involved in protein refolding, disaggregation, trafficking and degradation (Balchin et al., 2016). We reasoned that while the feeding mitotic peak (1dS) is a situation when *de novo* folding after translation is expected, starvation may be considered as a situation of stress needing cytosolic chaperones. Since DR in rodents and humans is known to increase the expression of the HSP70 family of chaperone proteins in skeletal muscle (Selsby et al., 2005; Yang et al., 2016), we wondered whether alternative analysis on the RNA-seq data could decipher changes in the expression also during 30 days of starvation. For that we grouped differentially expressed genes (DEGs) with similar expression profiles at 1dS, 7dS and 30dS into 6 clusters by fuzzy c-means clustering (Guthke et al., 2005) (Figure 1C and Table S2A). Interestingly, cluster 1 shows a U shape with a relative expression increase at 1dS and 30dS respect to 7dS, and “metabolic processes” and “protein folding” are the biological processes most enriched which include 16 out 26 of the cytosolic chaperones previously seen (Table S2B-C). Moreover, we found that most (23/27) of the transcripts related to protein folding in cluster 1 to be enriched in X1 (stem cells) when compared to Xins (differentiated cells) (Hayashi et al., 2006) (Table S2D). It drew our attention that a member of the proteasome (i.e. proteasome 26S subunit or PSMD5) as well as most of the subunits of the chaperonin TRiC/CCT are present in cluster 1 and enriched in stem cells (6/8) (Figure 1D and Table S2D). A search for components of the proteasome and the remaining CCT subunits not included in cluster 1 corroborated that they are enriched in stem cells, and at least the members of TRiC follow a trend of up-regulation at 1dS and 30dS compared to 7dS (Figure 1D, S1B and Table S2D). In agreement with our observation it has been suggested that increased proteasome and high levels of CCT subunits are hallmarks of hESCs (Noormohammadi et al., 2016; Vilchez et al., 2012a). However they may be involved in different functions since hESCs display high levels of protein synthesis, whereas we found that stem cell enriched translation is down-regulated at 7dS and 30dS when compared to 1dS (Figure 1A-B and Table S2D). TRiC seems to be a potential regulator of starved stem cells and studies in this direction are lacking. Therefore we decided to further characterize the function of TRiC specifically during starvation.

### TRiC is necessary for regeneration specifically in starved planarians

TRiC conforms a double-ring complex of 8 units codified by different genes (*cct-1* to *cct8*) that belong to the chaperone family of HSP60 (Kubota et al., 1995). Homologs of each *cct* gene have been previously identified in the sexual strain of *S. mediterranea* (*Smed-cct1A*, *Smed-cct2*, *Smed-cct3A*, *Smed-cct4B*, *Smed-cct5*, *Smed-cct6*, *Smed-cct7* and *Smed-cct8A*) which are expressed in the soma and testes (Counts et al., 2017). Paralogs for some of them (*Smed-cct1B*, *Smed-cct3B, Smed-cct4A* and *Smed-cct8B*) were also found to be exclusively expressed in the testes and specifically required for sperm elongation in the sexual strain (Counts et al., 2017; Rouhana et al., 2017). By feeding sexual planarians with RNAi of the somatic *ccts*, it was observed that they are required for homeostatic survival not linked to a decrease in stem cells (Counts et al., 2017). In the asexual strain the testes-specific *ccts* have an almost or even completely (i.e. *cct4A*) undetectable expression as seen in our RNA-seq data sets and the Dresden reference transcriptome (PlanMine) (Rozanski et al., 2019) and therefore we focus on the somatic *ccts*.

Aiming to understand the function of *ccts* in starved stem cells, we designed RNAi schedules to study the stem cells under different nutritional status. We injected planarians with either dsRNA of *gfp* as control or any of the *ccts* for three consecutive days and then amputated heads and tails 5 days later to follow the regeneration of the trunks. The induction of regeneration aimed to amplify the effect of the RNAi specifically in the stem cells, since amputation induces mitosis during regeneration (Baguñà, 1976b; Saló and Baguñà, 1984; Wenemoser and Reddien, 2010). In one of the schedules, the planarians were at 14 days of starvation when performing the amputation (Figure 1E). We independently down-regulated the 8 *ccts* which showed similar phenotypes consisting in planarians not being able to form a normal regenerating blastema (new regenerating tissue) and starting to die 20 days after amputation (Figure 1E and S1C). Remarkably when one extra feeding was allocated one day before the first injection and thus planarians were at 8dS when amputated, most of the planarians were able to form a blastema and fully regenerate (Figure 1F and S1D) even though the levels of mRNA were significantly down-regulated as checked for *cct3A* RNAi (Figure S1E). Next, we chose one of the *ccts* with the highest phenotype penetrance (*Smed-cct3A*) and performed a schedule of long-term starvation in which planarians were at 37 days of starvation when performing the amputation obtaining the same phenotype as the one in which they were 14 days starved (Figure S1F). Hereinafter the shortest schedule (14 days of starvation when performing the amputation) will be referred as “starved conditions” whereas the one with one feeding before the first injection will be referred as “feeding conditions”. We conclude that TRiC is necessary for blastema formation in starved but not in fed planarians.

### Down-regulation of *cct3A* in starved planarians causes an accumulation of mitoses immediately after both regenerative mitotic peaks eventually leading to stem cell exhaustion

The observation that *ccts* are necessary for blastema formation suggested that they may regulate stem cell proliferation and/or differentiation. Therefore we examined the pattern of mitoses during regeneration in starved planarians by using a Histone H3 phosphorylated at serine 10 antibody (anti-H3P) (Hendzel et al., 1997). It has been shown that planarian amputation triggers two mitotic peaks early in regeneration that contribute to blastema formation and growth (Baguñà, 1976b; Wenemoser and Reddien, 2010). We observed that *cct3A(RNAi)* animals had an increased number of H3P^+^ stem cells at the mitotic minimum that happens just after the first mitotic peak at 12 and 20 hours of regeneration (12hR and 20hR) and also just after the second mitotic peak (48hR, 72hR and 4dR) when compared to control planarians (Figure 2A-B). At 15dR *cct3A(RNAi)* animals also showed a slight increase of mitoses, whereas after 50dS mitotic activity was nearly abolished when compared to controls (Figure 2A-B).

**Figure 2.**
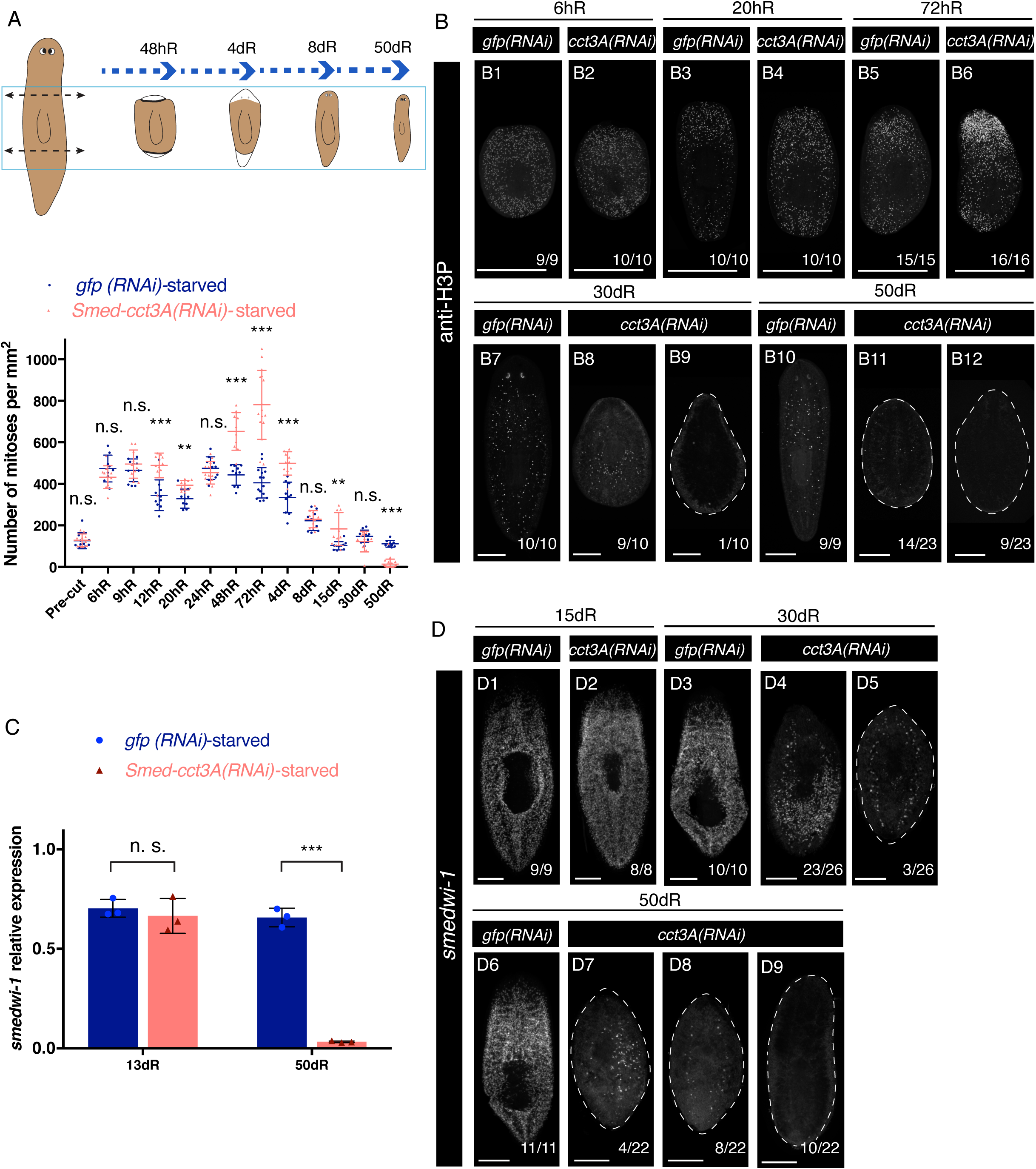
*cct3A* RNAi shows increased numbers of mitoses after both mitotic peaks and stem cell depletion at late time points of regeneration. (A) The schematic displays the double amputation (double arrows) performed on the RNAi injected planarians. The trunks are kept to follow regeneration. The graph shows the mitotic numbers during different time points of regeneration. Error bars are s.d. from the mean and asterisks indicate P < 0.01 (two asterisks), P < 0.001 (three asterisks) and n.s. indicates not significant using two-tailed Student’s test with equal variance. n ≥ 9 planarians per time point. (B) Maximum projections of representative trunks labelled with anti-H3P at different time points of regeneration. On the bottom, the number of planarians with the phenotype shown from the total is displayed. At 20hR (B4) there is an increased number of mitoses in *cct3A(RNAi)* planarians which accumulate at the wound site at 72hR (B6) compared to controls. At 30dR and 50dS there are some planarians with no mitoses (B9 and B12). (C) Relative expression of *smedwi-1* at 13 and 50 days of regeneration after either *cct3A* or *gfp* RNAi. Error bars are s.d. from the mean. Asterisks indicate P < 0.001 (three asterisks) and n.s. indicates not significant using two-tailed Student’s test with equal sample variance. n = 3 replicates (5 planarians each) per time point. (D) Maximum projections of representative trunks after FISH for *smedwi-1* at different time points of regeneration. On the bottom, the number of planarians with the phenotype shown from the total is displayed. While at 15dR *cct3A(RNAi)* planarians display similar expression of s*medwi-1* as controls (D2), at 30dR some planarians have almost no expression (D5) and at 50dR almost all planarians show no expression (D8 and D9). hR, hours of regeneration; dR, days of regeneration. Scales, 1 mm (B1-B6) and 250 µm (B7-B12 and D1-D9).

Using *smedwi-1*, a marker that labels most proliferating planarian adult stem cells (Reddien et al., 2005), for fluorescent *in situ* hybridizadion (FISH) and for qPCR (Figure 2C-D) we sought to determine whether *cct3A* RNAi affects stem cell numbers during starvation. At 13dR most of the treated planarians displayed a normal distribution and levels of *smedwi-1* (Figure 2C-D), but at 30dR stem cell loss was evident in some planarians and by 50dR, most of the planarians showed very few if any stem cells (Figure 2C-D). In agreement with the lack of blastema growth in *cct3A(RNAi)* planarians (Figure 1E), we observed minimal differentiation of eyes (anti-VC1) (Sakai et al., 2000), brain (*Smed-gpas*) (Iglesias et al., 2011), epidermal cilia (anti-acTUB) (Reddien et al., 2007; Robb and Sanchez Alvarado, 2002) and muscle (anti-TMUS) (Cebrià et al., 1997) in anterior wounds (Figure S2A).

As expected, this effect on mitosis and total number of stem cells is specific of starvation since *cct3A* RNAi on feeding conditions does not affect the number of mitoses at 72hR (Figure S2B) and planarians show normal distribution and expression of *smedwi-1* at 30dR (Figure S2C-D) in agreement with the already observed normal regeneration phenotype (Figure 1F).

### Mitotic failure, genome instability, increased vacuolization and cell death are induced at the wound site after *cct3A* down-regulation during starvation

Since the increase of mitotic cells observed in *cct3A(RNAi)* planarians does not translate into blastema overgrowth or tumour formation like after down-regulation of certain tumour suppressors in planarians (Gonzalez-Estevez et al., 2012b) and differentiation is minimal, we suspected that mitotic stem cells were dying. Therefore we performed TUNEL staining at 72hR, as the time point when the difference in mitosis is the highest in respect to controls and at 4dR when the difference decreases again (Figure 2A). Although the differences in the number of TUNEL^+^ cells at the wound site were not significantly different between *cct3A(RNAi)* planarians and controls at 72hR, at 4dR a massive and significant increase of TUNEL^+^ cells was observed in both anterior and posterior wounds (Figure 3A).

**Figure 3.**
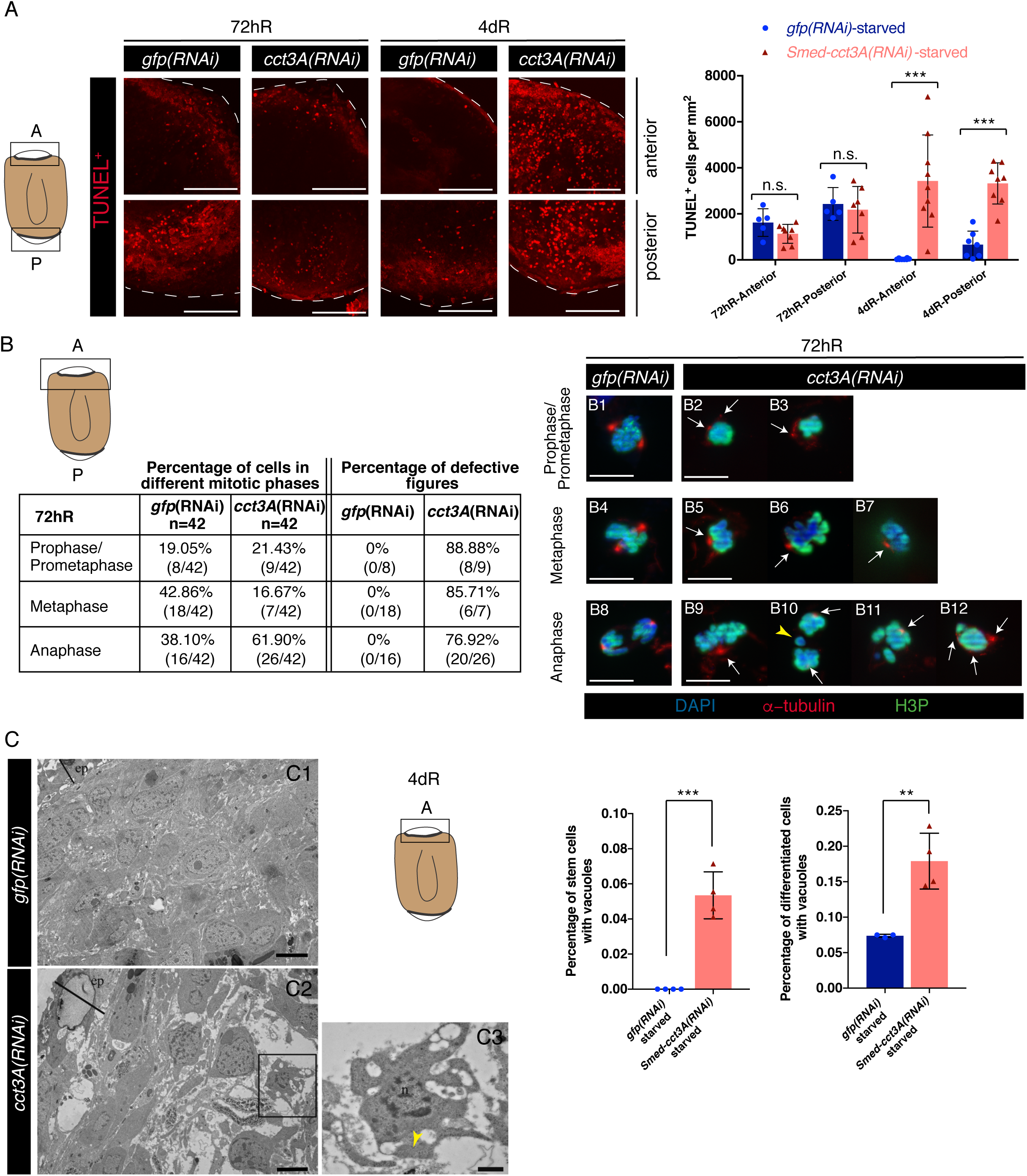
*cct3A* RNAi during starvation leads to mitotic defects, chromosomal aberrations, cell death and vacuolization at wound sites early during regeneration. (A) The cartoon represents a regenerating trunk and the squares display the region analyzed at the anterior “A” and posterior “P” blastemas. The images are TUNEL maximum projections of representative blastemas from 72 hours and 4 days regenerating trunks. The graph shows the number of TUNEL^+^ cells per mm^2^. The differences in cell death are not significant (n.s.) at 72hR (two-tailed Student’s test with equal sample variance) and significant (***P < 0.001 using two-tailed Student’s test with equal variance) at 4dR. n ≥ 5. (B) The cartoon represents a regenerating trunk and the square displays the region analyzed at the anterior “A” blastema. Quantification of the percentage of stem cells in different mitotic phases and the percentage of defective mitotic figures in 72hR anterior blastemas of *cct3A* RNAi and controls after double immunostaining with anti-α-tubulin and anti-H3P. Representative images are shown. Nuclei are stained with DAPI. Arrows indicate abnormal organization or number of spindle poles and the yellow arrowhead indicates chromosome lagging. (C) Micrographs display the wound region of *cct3A* RNAi and controls at 4dR. *cct3A(RNAi)* planarians show reduced cell density and cells with vacuoles. The cartoon represents a regenerating trunk and the square displays the region analyzed. The box in C2 indicates the high magnification image (C3) which shows a differentiated cell with vacuoles in the cytoplasm. Quantification of differentiated and stem cells with vacuoles in the blastema region shows that *cct3A* RNAi leads to increased vacuolization in differentiated and stem cells. Each value is the average ± standard deviation of three independent samples. For each sample, more than 50 blastemal cells were counted. Asterisks indicate P < 0.01 (two asterisks) and P < 0.001 (three asterisks) using two-tailed Student’s test with equal sample variance. Arrow head indicates mitochondria with normal morphology. dR, days of regeneration; ep, epithelium; n, nucleus. Scales, 4 µm (A), 10 µm (B), 4 µm (C1 and C2) and 1 µm (C3).

We speculated that the mitotic stem cells that accumulate at 72hR are dying at 4dR because of a hampered progression through the different phases of mitosis. We examined mitotic stem cells from 72hR *cct3A(RNAi)* wounds by double immunostaining with anti-*α*-tubulin (de Sousa et al., 2018) and anti-H3P. Significantly, we observed a decrease in the percentage of stem cells in metaphase (from ∼ 43% in controls to ∼17%) and an increase in the percentage of cells in anaphase (from ∼38% in controls to ∼62%) (Figure 3B), suggesting that mitotic cells get stuck in anaphase at 72hR. In addition, more than 70% of all mitotic figures are defective (Figure 3B). The alterations included abnormal organization or number of spindle poles, in agreement with the role of TRiC in folding tubulin (reviewed in (Spiess et al., 2004), asymmetrical karyokinesis (Figure 3B11 and B12) and chromosome lagging (Figure 3B10), overall leading to gross nuclear alterations. Electron micrographs also showed a reduced cellular density at the wound of 4dR *cct3A(RNAi)* animals compared to controls (Figure 3C). Unexpectedly we also observed that a significantly high number of both stem cells and differentiated cells displayed cytoplasm vacuolization when compared to controls (Figure 3C). The vacuoles were not double-membrane vesicles thus ruling out autophagy as the involved cellular process. However they seem to be internalizing extracellular fluid or small content, thus suggesting that *cct3A* RNAi increases pinocytosis, a form of endocytosis involving fluids and solutes which in humans is linked to the absorption of fat droplets in the small intestine (Stillwell, 2016), antigen presentation in dendritic cells and macrophages and to acquire nutrients from the environment in cancer cells (Commisso et al., 2013).

Altogether, our data suggest that *cct3A* is essential to allow mitosis in starved regenerating animals.

### *cct3A* down-regulation induces a transcriptional compensatory mechanism and down-regulates the UPR^ER^ during regeneration of starved planarians

In order to understand the signalling pathways that are affected by under the control of *cct3A*, we performed RNA-seq of whole starved *cct3A(RNAi)* planarians and controls at 72hR. The differential analysis found up-regulated transcripts that belong to the GO category of ‘organonitrogen compound catabolic process’ and ‘alpha-amino acid metabolic process’ suggesting that *cct3A(RNAi)* planarians are increasing these metabolic pathways presumably to obtain energy at 72hR (Figure 4A and Table S3A). The other enriched biological process enriched was protein folding where most of the transcripts are *ccts* or other chaperones (Figure 4A-B and Table S3B). The remaining transcripts in this category include several tubulin and actin transcripts which are two of the known clients of TRiC (reviewed in (Spiess et al., 2004) and some genes known to enhance general protein folding and assist with the folding of tubulin (Table S3B). This altogether suggests that upon down-regulation of *cct3A* a compensatory mechanism is activated that tries to counteract for the decreased protein folding capacity.

**Figure 4.**
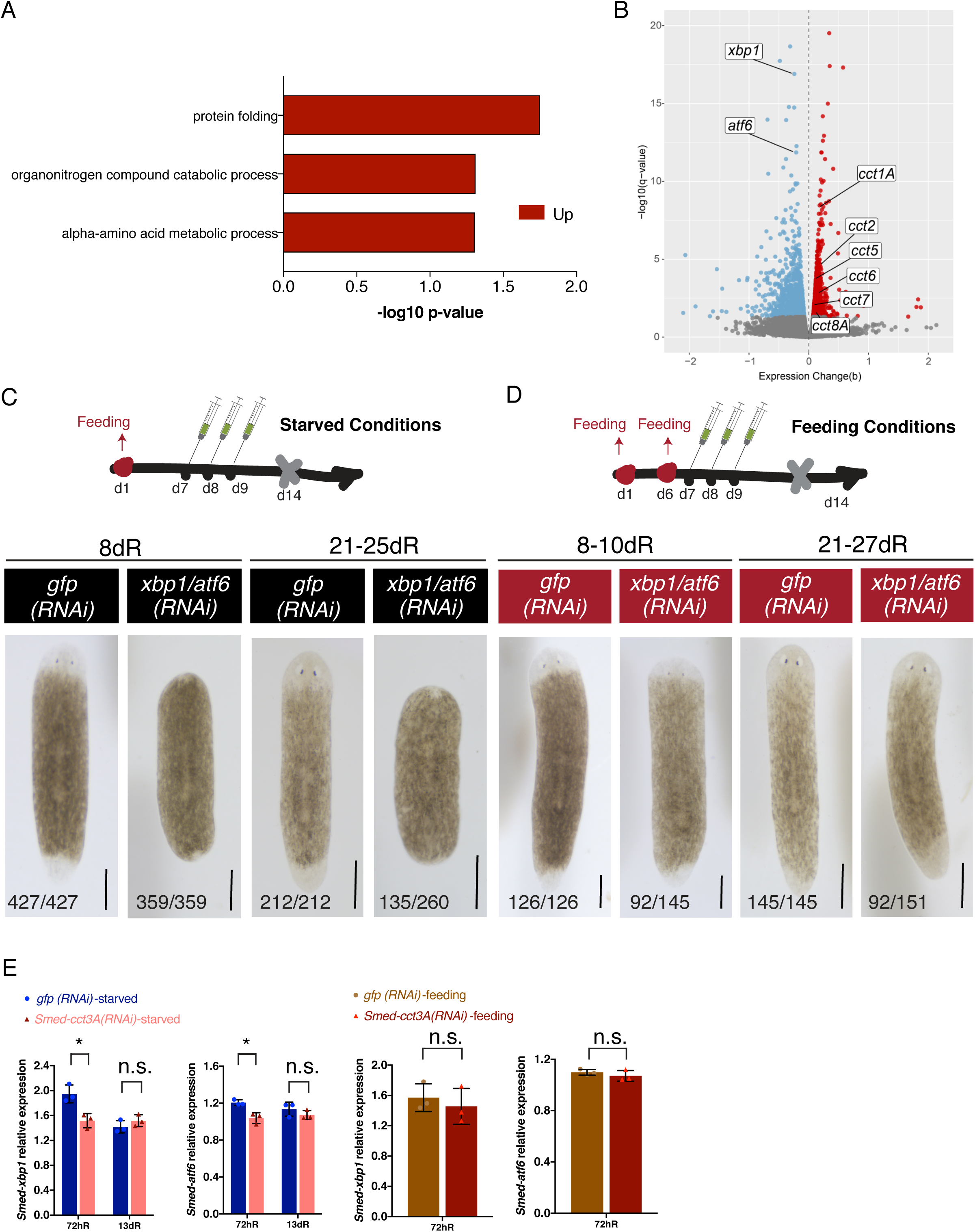
*cct3A* RNAi down-regulates *xbp1* and *atf6* at 72hR specifically during starvation and double RNAi for both genes leads to a similar phenotype as *cct3A* RNAi. (A) GO enrichment analysis for biological processes of the 982 up-regulated genes (q-value < 0.05) in *cct3A* RNAi planarians versus controls at 72hR. No enriched biological processes were found down-regulated. For better visualization, similar enriched GO terms based on the same subset of genes were manually removed to reduce redundancy. (B) Volcano plot displaying DEGs in *cct3A(RNAi)* planarians compared to controls at 72hR (q-value < 0.05). Blue and red dots represent down- and up-regulated transcripts, respectively. Y-axis indicates the negative log10 of the false discovery rate (FDR) (q-value). X-axis indicates the beta values (b) which is a biased estimator of expression change (fold change) given by Sleuth. For better visualization, dots with -log10 (q-value) > 20 and |(b)| > 2.2 are not displayed. (C) RNAi injections schedule in starved conditions. Planarians are at 14dS when the amputation is performed (indicated by the grey cross at day 14). Live images show that *xbp1/atf6(RNAi)* planarians form a minimal blastema when compared to controls at the time points shown. At the bottom are the number of planarians with the phenotype shown. The remaining planarians are dead by the time point of regeneration shown. (D) RNAi injections schedule in feeding conditions. One extra feeding respect to C is introduced one day prior to injections. Planarians are at 8dS when the amputation is performed (indicated by the grey cross at day 14). The live images show that most of the *xbp1/atf6(RNAi)* planarians regenerate as controls. At the bottom are the number of planarians with the phenotype shown. The remaining planarians at 8-10dR are either dead (13/145) or have a tiny blastema and died later (40/145). The remaining planarians at 21-27dR are dead (E) Relative expression of *xbp1 and atf6* at 72 hours and 13 days of regeneration after either *cct3A* or *gfp* RNAi during either starving or feeding conditions. The graphs show that *cct3A* RNAi down-regulates *xbp1* and *atf6* specifically during starvation at 72hR. Error bars are s.d. from the mean. Asterisks indicate P < 0.05 (one asterisk) and n.s. indicates not significant using two-tailed Student’s test with equal sample variance. n = 3 replicates (5 planarians each) per time point. dR, days of regeneration; hR, hours of regeneration. Scales, 500 µm.

Validating our previous results, we found genes related to genome instability and to endocytosis among the up-regulated transcripts (Table S3C-D). Strikingly, the top down-regulated genes upon *cct3A* RNAi are the components of the unfolded protein response on the endoplasmic reticulum (UPR^ER^) *atf6* and *xbp*1 (Figure 4B). These are two transcription factors that are crucial for two of the three main UPR^ER^ branches responsible for the recovery of ER homeostasis or the induction of apoptosis (Labbadia and Morimoto, 2015). We conclude that under starvation the cytosolic chaperone *cct3A* induces the UPR in the ER suggesting a crosstalk regulation between the cytosol and the ER.

### TRiC induces the unfolded protein response, which is essential for mitosis progression during regeneration of starved planarians

Our observations raise the question whether the phenotype observed after *cct3A* down-regulation is due to ER stress and the inability to resolve this stress by the UPR^ER^. In order to investigate this aspect, we performed RNAi experiments for *Smed-xpb1* and *Smed-atf6*. Although individual RNAi for either *xbp1* or *atf6* did not give any apparent phenotype after two rounds of injections, double RNAi *xbp1/atf6* (Figure 4C-D) displayed the *cct3A(RNAi)* phenotype of no regeneration in starved conditions and full regeneration under feeding conditions, albeit with a lower percentage of feeding rescue than in *cct3A(RNAi)* (∼61% in *xbp1/atf6 (RNAi)* compared to ∼84% in *cct3A(RNAi)* (Figure 1F and 4D). The level of down-regulation of *xbp1* and *atf6* in the double RNAi was significantly different between controls and treated planarians in the starved and the feeding condition (Figure S3A). We validated by qPCR the decreased expression of *xbp1* and *atf6* upon *cct3A* RNAi at 72hR (Figure 4E). Interestingly we also found that the down-regulation of UPR^ER^ regulators in *cct3A* RNAi is specific of starvation (Figure 4E and F) and early regeneration (Figure 4E) since it is not maintained at later time points of regeneration (13dR) (Figure 4E). Furthermore, *xbp1* expression is linked to starvation (upregulated at 7dS and 30dS respect to 1dS; cluster 3 in Figure 1C) (Figure S3B and Table S2A).

Next we checked the mitotic profile of regenerating *xbp1/atf6(RNAi)* planarians compared to controls during starvation. Although *xpb1/atf6(RNAi)* planarians are able to mount a first mitotic response at 6hR, they do not show a second mitotic response since the number of mitoses is significantly lower than in controls at 48hR and 72hR (Figure 5A) compared to the feeding condition which shows a normal mitotic index for the same time points (Figure S4A). Whereas *cct3A(RNAi)* animals died because of stem cell exhaustion, most of the *xbp1/atf6(RNAi)* planarians had normal levels of mitosis and stem cells before dying (Figure5A-C) with only some planarians showing no mitosis and no stem cells between 15dR and 23dR (Figure 5A5, C3-C4). This agrees with both *xbp1* and *atf6* being enriched in X2 (differentiating cells) and Xins (differentiated cells) in addition to be expressed in stem cells (Figure S4B).

**Figure 5.**
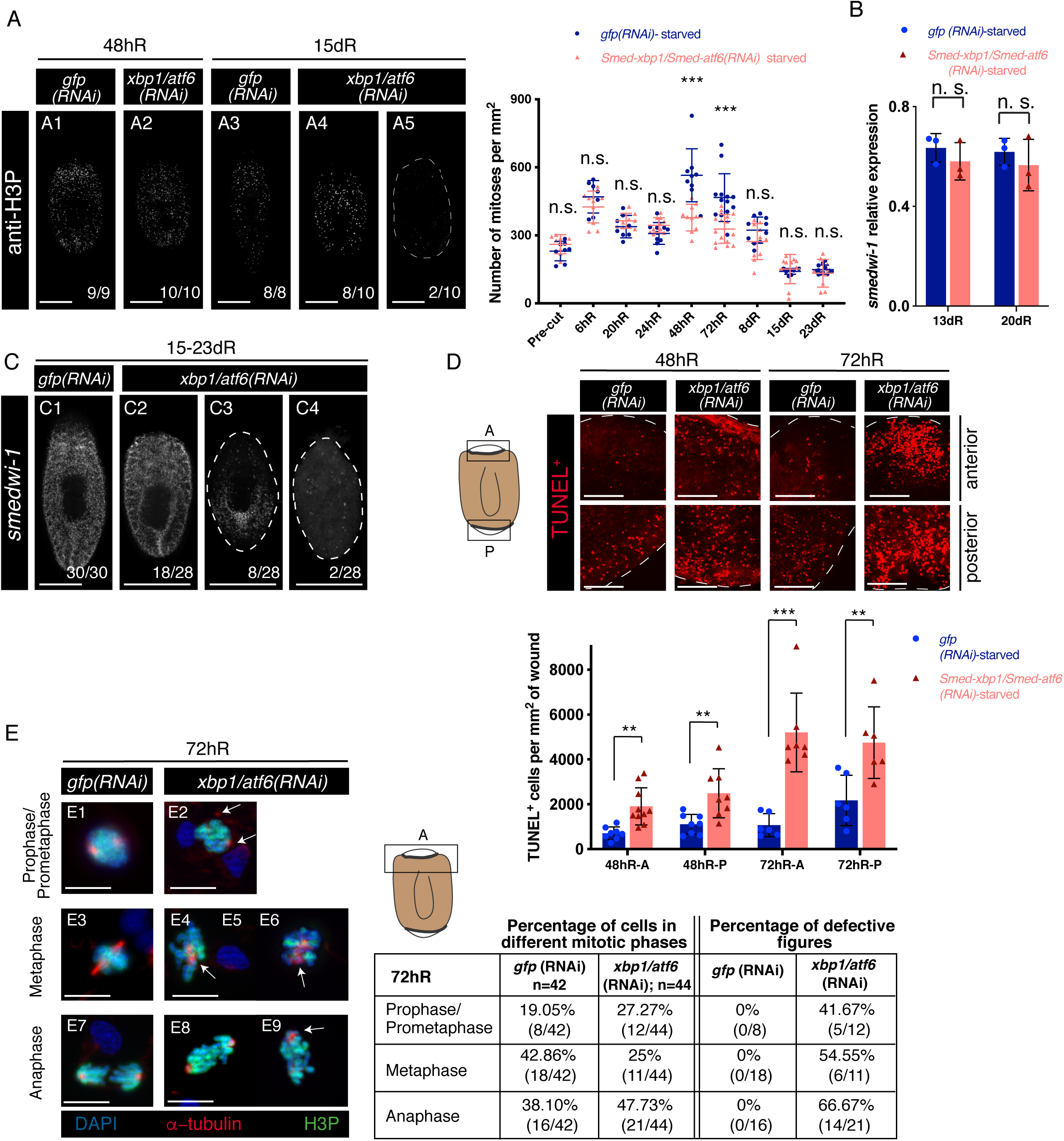
*xbp1/atf6* RNAi displays most of the phenotypes of *cct3(RNAi)* planarians. (A) Maximum projections of representative trunks labelled with anti-H3P at different time points of regeneration. On the bottom, the number of planarians with the phenotype shown from the total is displayed. At 48hR there is a decreased number of mitoses in *xbp1/atf6(RNAi)* planarians (A2) compared to controls. At 15dR some *xbp1/atfg6(RNAi)* planarians display almost no mitoses (A5). The graph shows mitotic numbers during different time points of regeneration. Error bars are s.d. from the mean and asterisks indicate P < 0.001 (three asterisks) and n.s. indicates not significant using two-tailed Student’s test with equal variance; n ≥ 7 planarians per time point. (B) Relative expression of *smedwi-1* at 13 and 20 days of regeneration after either *xpb1/atf6* or *gfp* RNAi. Error bars are s.d. from the mean. n.s. indicates not significant using two-tailed Student’s test with equal sample variance. n = 3 replicates (5 planarians each) per time point. (C) Maximum projections of representative trunks after FISH for *smedwi-1* at 15-23 days of regeneration. On the bottom, the number of planarians with the phenotype shown from the total is displayed. Although the level of *smedwi-1* expression is similar between *xbp1/atf6* RNAi and controls, some planarians display almost no expression (C3-C4). (D) The cartoon represents a regenerating trunk and the squares display the region analyzed at the anterior “A” and posterior “P” blastemas. The images are TUNEL maximum projections of representative blastemas from 48 and 72 hours regenerating trunks. The graph shows the number of TUNEL^+^ cells per mm^2^ at the same time points in anterior and posterior blastemas. Error bars are s.d. from the mean and asterisks indicate P < 0.01 (two asterisks) and P < 0.001 (three asterisks) using two-tailed Student’s test with equal variance. n ≥ 6 planarians per time point. Cell death is increased in *xbp1/atf6(RNAi)* blastemas at both time points. (E) Representative images of stem cells from 72 hours regenerating blastemas labelled with anti-α-tubulin, anti-H3P and DAPI. Arrows indicate abnormal organization or number of spindle poles. E8 displays asymmetrical distribution of chromosome content. The cartoon represents a regenerating trunk and the square displays the region analyzed at the anterior “A” blastema. The table displays the quantification of the percentage of stem cells in different mitotic phases and the percentage of defective mitotic figures. Note that the *gfp* controls are the same as displayed in Figure 3B, since the experiment was performed at the same time with the same controls for both conditions. dR, days of regeneration; hR, hours of regeneration. Scales, 500 µm (A and C), 100 µm (D) and 10 µm (E).

In line with the lack of blastema growth in *xbp1/atf6(RNAi)* planarians similar to *cct3A* RNAi (Figure 1E), we observed minimal differentiation of eyes (anti-VC1) (Sakai et al., 2000), brain (*Smed-gpas*) (Iglesias et al., 2011), epidermal cilia (anti-acTUB) (Reddien et al., 2007; Robb and Sanchez Alvarado, 2002) and muscle (anti-TMUS) (Cebrià et al., 1997) in anterior wounds (Figure S4C).

Comparable to *cct3A* RNAi at 4dR, cell death increases at 48hR and 72hR in both anterior and posterior *xbp1/atf6(RNAi)* wounds as detected by TUNEL (Figure 5D). This observation, together with the low number of mitoses observed at the same time point, suggests that stem cells are dying during mitosis. A closer look at the mitoses at 72hR showed a similar phenotype as in *cct3A* RNAi with a high percentage of defective figures (between ∼42% and ∼67% depending on the mitotic phase) including mislocalized or abnormal number of spindle poles (Figure 5E) and asymmetrical karyokinesis (Figure 5E8) but chromosome lagging was not observed. Significantly, we observed a decrease in the percentage of stem cells in metaphase (from ∼43% in controls to ∼25%) similar to that observed upon *cct3A* RNAi and only a slight increase in the percentage of cells in anaphase (from ∼38% in controls to ∼48%) (Figure 5E), suggesting that mitotic cells are not stuck at anaphase as much as in *cct3A(RNAi)* animals.

Our data strongly suggests that most of the effects of *cct3A* down-regulation are due to down-regulation of the UPR.

### During starvation *cct3A(RNAi)* and *xbp1/atf6(RNAi)* planarians show defects in lipid droplet metabolism with *cct3A* RNAi resulting in decreased energy stores

We suspected that the dependency of TRiC and UPR phenotypes to starvation rely on affecting certain metabolic processes. Supporting evidence is the up-regulation of amino acid metabolism and organonitrogen compound catabolism seen upon *cct3A* RNAi (Figure 4A). In order to investigate this point further we performed RNA-seq of 72hR *xbp1/atf6(RNAi)* planarians. By GO enrichment analysis (Figure 6A and Table S4A) we observed an increase in apoptosis supporting our data on TUNEL staining. *xpb1/atf6* RNAi also seems to increase translation (GOs “tRNA metabolic process” and “amino acid activation”) (Figure 6A), which is known to be down-regulated upon activation of the UPR in other organisms (Labbadia and Morimoto, 2015). Interestingly, apart from the expected general down-regulation of the response to ER stress or protein folding, the remaining down-regulated GO-terms referred to metabolic processes including lipid metabolism (Figure 6A, Table S4B). Remarkably, a closer look into the RNA-seq of *cct3A* RNAi, also retrieved a repertoire of DEGs related to lipid metabolism (Figure 6B). Among them we found DEGs related to lipid droplet (LD) biogenesis (Pol et al., 2014) with most of them upregulated in *cct3* RNAi (12/16) while down-regulated in *xbp1/atf6* RNAi (29/30) (Figure 6B and Table S5A). LDs are intracellular organelles specialized to store fatty acids (FAs) and to channel them into pleiotropic pathways including phospholipid biosynthesis, energy production via β-oxidation and TCA cycle, lipoprotein generation and the formation of bioactive lipid mediators. LD formation requires the coordination of several processes: (1) fatty acid activation, (2) synthesis of neutral lipids, (3) remodelling of phospholipids, (4) synthesis of new phospholipids, and (5) the function of accessory proteins (Pol et al., 2014). Indeed we found key transcripts for these sub-processes of LD biogenesis to be up-regulated in *cct3A(RNAi)* animals and down-regulated in *xbp1/atf6(RNAi)* animals. For instance, we found acyl-CoA synthetase which activates fatty acids, ethanolamine phosphate cytidyltransferase which provides CDP-ethanolamine required for the biosynthesis of phosphatidylethanolamine (a major phospholipid) or perilipins which are an excellent example for an accessory protein (Figure 6D and Table S5A). Sphingolipids are part of the phospholipid monolayer of LDs. We also found the biological process “sphingolipid biosynthetic process” with representative genes up-regulated in *cct3A* RNAi (3/5) (Figure 6B) and most of them down-regulated in *xbp1/atf6* RNAi (5/7) (Table S5A). Altogether our data suggests that *cct3* and *xbp1/atf6* RNAi have an antagonistic role in LD biogenesis.

**Figure 6.**
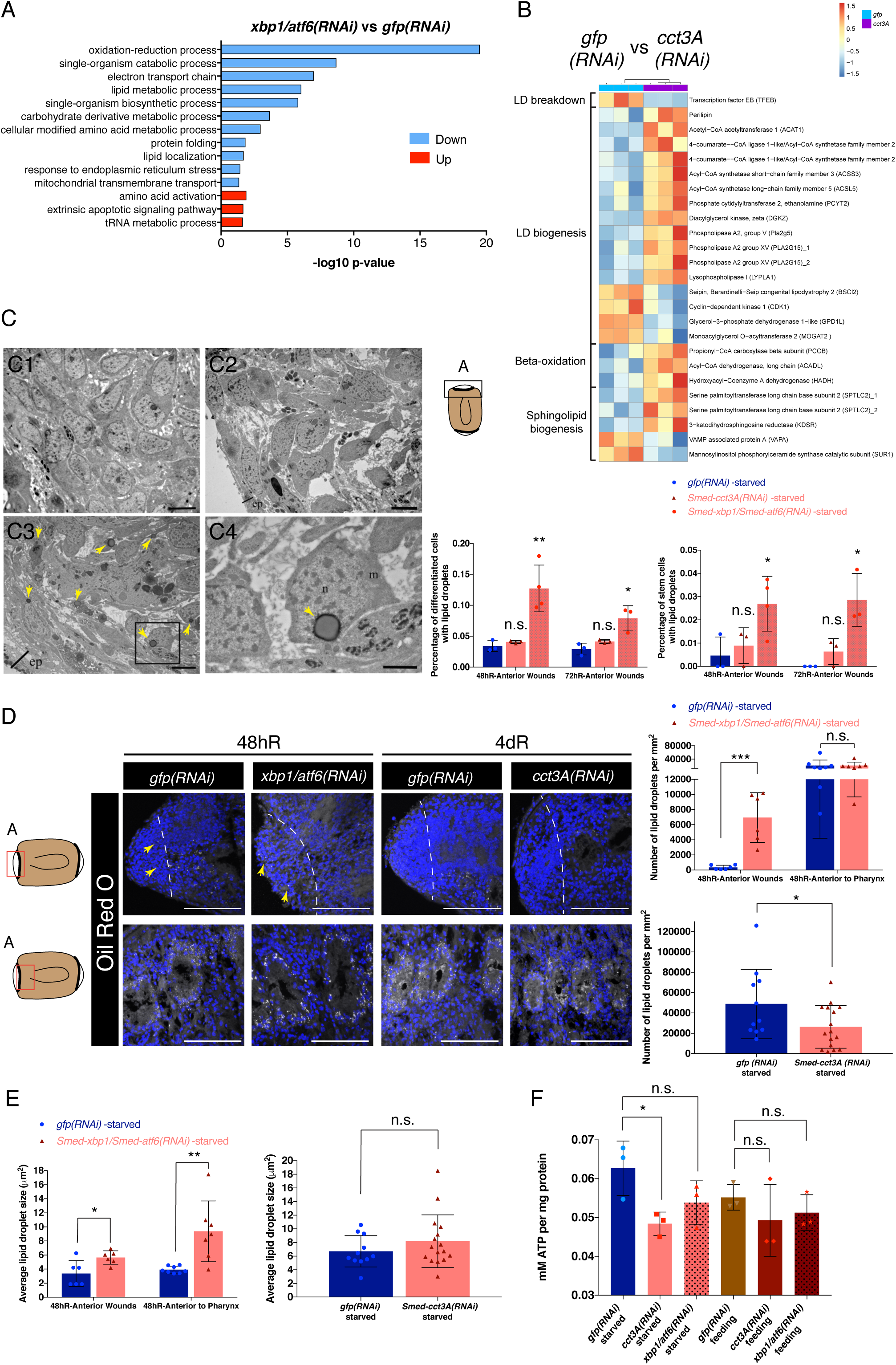
*xpb1/atf6* and *cct3A* RNAi modulate lipid metabolism with *cct3A* RNAi leading to decreased energy storage. (A) GO enrichment analysis for biological processes of the 1431 up-regulated genes (bars in red) and the 1735 down-regulated genes (bars in blue) (q-value < 0.05) in *xbp1/atf6*(RNAi) planarians versus controls at 72hR. For better visualization, similar enriched GO terms based on the same subset of genes were manually removed to reduce redundancy. (B) The heat map shows the expression of DEGs in *cct3A* RNAi at 72hR involved in different process related to LD metabolism. The colour-coded scale covers the range of TPM (transcripts per million) values per replicate. (C) Analysis of *cct3A(RNAi)* and *xbp1/atf6(RNAi)* planarian wounds at 48 hours and 72 hours regeneration by TEM. C1 corresponds to *gfp* RNAi, C2 to *cct3A* RNAi showing less cell density than controls and C3 corresponds to *xbp1/atf6* RNAi showing less cell density and more cells with lipid droplets (LDs) (arrow heads) than in controls. The square indicates the high magnification shown in C4, a stem cell with a LD. The cartoon represents a regenerating trunk and the square displays the region analyzed at the anterior “A” blastema. The graphs show the quantification of differentiated and stem cells with LDs in anterior wounds at 48 hours and 72 hours of regeneration. Each value is the average ± standard deviation of three independent samples. For each sample, more than 50 blastemal cells were counted. Asterisks indicate P < 0.05 (one asterisks) and P < 0.01 (two asterisks) and n.s. indicates not significant respect the corresponding controls using two-tailed Student’s test with equal sample variance. *xpb1/atf6* RNAi have more differentiated and stem cells with LDs than controls at both time points. ep, epithelium; m, mitochondria; n, nucleus. (D) The cartoon represents a regenerating trunk and the squares display the region analyzed at the anterior “A” region. Panels are representative maximum projections of cryosections stained with Oil Red O to label LDs. Arrow heads indicate LDs. The first graph shows the quantification of LDs in either the wound site or a just anterior to the pharynx in *xbp1/atf6* RNAi at 48 hours of regeneration showing an increased number specifically at wound sites in *xbp1/atf6* RNAi. The second graph is the quantification of LDs anterior to the pharynx in *cct3A* RNAi at 4 days of regeneration showing a decreased number compared to controls. At the wound site at 4 dR there are no LDs for any of the conditions analyzed. Error bars are s.d. from the mean and asterisks indicate P < 0.05 (one asterisk), P < 0.001 (three asterisks) and n.s. indicates not significant using two-tailed Student’s test with equal variance. n ≥ 6 planarians per time point. (E) The graphs correspond to the data of graphs in D, displaying the average LD size which is higher than controls for both regions in *xpb1/atf6* RNAi. Error bars are s.d. from the mean and asterisks indicate P < 0.05 (one asterisk), P < 0.01 (two asterisks) and n.s. indicates not significant using two-tailed Student’s test with equal variance. (F) ATP measurement on 48hR trunk extracts shows that *cct3A* RNAi under starved conditions contains less ATP than controls. Error bars are s.d. from the mean and asterisks indicate P < 0.05 (one asterisk) and n.s. indicates not significant using two-tailed Student’s test with equal variance. n = 3 replicates (5 planarians per replicate). dR, days of regeneration; hR, hours of regeneration. Scales, 4 µm (C1-C3), 1 µm (C4) and 100 µm (D).

Cells break down neutral lipids from the LDs when there is need of energy in the form of fatty acids and/or when membranes need to be synthesized. In mammals, the breakdown is done by the step-wise action of three main lipases: the adipose tissue lipase (ATGL), the hormone-sensitive lipase (HSL) and the monoglyceride lipase (MGL). Remarkably *xbp1/atf6* RNAi shows down-regulation for ATGL and two MGL transcripts (Table S5B). While *cct3(RNAi)* animals do not differentially express those transcripts, they show a homolog for TFEB down-regulated (Figure 6B and Table S5A), a master regulator of lysosome biogenesis and autophagy, which is known to induce lipid breakdown in the mouse liver upon starvation (Settembre et al., 2013). Interestingly “fatty acid β-oxidation”, the process to breakdown fatty acids to obtain energy has representative genes up-regulated in *cct3A(RNAi)* (3/3) (Figure 6B) with most of them down-regulated in *xbp1/atf6(RNAi)* animals (15/15) (Table S5A).

Thus in *cct3A(RNAi)* planarians, LD biosynthesis and β-oxidation of fatty acids are up-regulated while LD breakdown is down-regulated. In the case of *xbp1/atf6(RNAi)* planarians, LD biogenesis and breakdown are down-regulated. Based on the RNA-seq data of both phenotypes *cct3A(RNAi)* and *xbp1/atf6(RNAi)* we hypothesized that LD homeostasis is disturbed in RNAi planarians. In order to investigate this further, we analyzed the number of cells with LDs at 48hR and 72hR blastemas by transmission electron microscopy (TEM) in *cct3A*(RNAi) and *xbp1/atf6(RNAi)* planarians (Figure 6C). Surprisingly, *cct3A* RNAi had a similar percentage of differentiated and stem cells with LDs as controls. However *xbp1/atf6* RNAi displayed a higher number (Figure 6C). In order to support our results on TEM, we performed Oil Red O staining on tissue sections to visualize the fat stored in LDs in blastema cells and also gastrodermal cells of digestive branches located in regions anterior to the pharynx (Figure 6D). In *xbp1/atf6(RNAi)* planarians we could corroborate the accumulation of cells with LDs in 48hR blastema cells and observed that this was specific of the blastema since gastrodermal cells displayed a similar number as controls (Figure 6D). Remarkably, in both blastema and gastrodermal cells, the average LD size is bigger than in controls (Figure 6E). Since we did not observe differences in LDs in *cct3(RNAi)* blastemas at 48hR and 72hR by TEM, we performed Oil Red O staining in anterior wounds at a later time point (4dR), when we previously observed an increase in stem cells and differentiated cells with vacuoles and increased cell death (Figure 6D). Although at the wound site we did not observe cells with LDs, in gastrodermal cells we observed a decreased number however bigger in size than controls (Figure 6E). In the case of *cct3A* RNAi, although LD biosynthesis, β-oxidation of fatty acids and sphingolipid biosynthesis are up-regulated, LD breakdown is down-regulated and there are less LDs in intestinal branches. This suggests that *cct3(RNAi)* animals are consuming the LDs either for energy production and/or anabolic processes without satisfying their requirements, which would explain why LD biogenesis is ongoing. The accumulation of LDs in *xbp1/atf6(RNAi)* planarians could be explained by either an increase in LD biogenesis or a decrease in LD mobilization. Our RNA-seq data favours the idea of LD mobilization being compromised since both LD biogenesis and breakdown of LDs are down-regulated. In order to understand whether RNAi planarians were unable to obtain energy from LDs, we determined general ATP levels in whole planarians (Figure 6F). We observed that control starved animals have surprisingly high ATP levels as controls that were fed, which hints towards a higher requirement to cope with the challenge for starved animals. Remarkably, *cct3(RNAi)* animals showed decreased levels of ATP specifically in starved conditions, thus confirming that *cct3(RNAi)* planarians do not obtain enough energy (Figure 6F) which may hamper homeostatic mechanisms thereby leading to enhanced stress and diminished regeneration. The levels of ATP were as controls during feeding and starved conditions in *xbp1/atf6*(RNAi) animals (Figure 6F), indicating that planarians are mobilizing LDs at some extent or/and that the local differences in energy are not detectable by our ATP assay.

## Discussion

Fasting protects stem cells and increases stem cell functionality through mechanisms which are not fully understood. Our knowledge of the mechanisms are limited to a few signalling pathways (Mihaylova et al., 2014): the regulation of the phosphatidylinositol 3-kinase/AKT (PI3K/AKT) pathway and the specific axis PKA-AMPK-EGR1 (Cheng et al., 2014; Di Biase et al., 2017) by insulin, the mechanistic target of rapamycin (mTOR) pathway by nutrients, the AMPK/LKB pathway by energy, and the Sirtuins through NAD^+^ levels. Although these pathways are potential important regulators of planarian stem cells during starvation, only mTOR has been previously characterized in that context (Iglesias et al., 2019). Here we found components of all these signalling pathways to be up-regulated in stem cells during planarian starvation and, most remarkably, we found a novel mechanism through which fasting regulates stem cells: the HSP60/chaperonin member TRiC (CCT, TCP1-ring complex or chaperonin containing TCP1) protects starved somatic stem cells, activates the UPR and allows planarian regeneration, specifically the mitotic response to amputation during starvation. We also demonstrate that RNAi for one of the subunits of TRiC, *cct3A*, leads to stem cell exhaustion, a hallmark of aging, only during starvation. Our findings also draw attention to the possibility of different regenerative programmes depending on nutritional status in many other organisms.

Growing evidence indicates that a tight regulation of proteostasis plays a major role in somatic stem cell self-renewal, pluripotency, and cell fate decisions. For instance during asymmetric cell divisions of NSCs, old or damaged proteins are channelled to progenitors in order to protect the stem cell (Moore et al., 2015). Similarly, high proteasome activity is required for ESC function since minimal down-regulation results in decreased levels of pluripotency markers and differentiation (Assou et al., 2009; Vilchez et al., 2012a). Cumulative evidence suggests that UPR^ER^ may be also important for somatic stem cell function since, for instance, BiP, a chaperone of the ER is essential for viability of pluripotent cells (Luo et al., 2006). Autophagy can also function as a selective mechanism for protein degradation (Nixon, 2013). It is necessary for cell reprogramming (Wang et al., 2013) and to maintain proteostasis of satellite cells (Garcia-Prat et al., 2016). Furthermore, highly proliferative cells such as cancer cells (Saez and Vilchez, 2014) or ESCs (Noormohammadi et al., 2018; Prinsloo et al., 2009) express high levels of certain chaperones (Noormohammadi et al., 2018) and increased levels of at least HSP90 and the chaperonin TRiC are necessary to regulate pluripotency components, confer ESCs with enhanced stress tolerance, and maintain them in a self-renewal mode since its down-regulation promotes differentiation (Bradley et al., 2012; Noormohammadi et al., 2016). In line with previous knowledge, we found that planarian stem cells show increased expression of several chaperones including most subunits of TRiC and proteasome components when compared to differentiated cells. However we observed that RNAi for components of TRiC or UPR^ER^ affect stem cells and regeneration mainly during starvation which was never described before.

Our results suggest that TRiC and probably other chaperones have different roles depending on the nutritional status, which agrees with the increased levels of expression for TRiC subunits and many other chaperones that we observed at 1dS and at 30dS. In planarians at 1dS (one day post-feeding) stem cells are highly proliferative, responding to feeding in a similar way like ESCs which are highly proliferative stem cells growing in nutrient-rich media with both showing high rates of protein synthesis. Therefore, a tight control of proteostasis may be necessary for *de novo* folding of proteins after translation. By contrary at 30dS we observed reduced levels of protein synthesis and thus the activation of TRiC would correspond to a response to stress. Under mild stress (7dS), high levels of TRiC may not be needed and thus stem cells may be able to overpass or avoid major problems after down-regulation of TRiC subunits. This last statement is in line with a report on *Drosophila* intestinal stem cells (ISCs). The authors challenged proteostasis by blocking components of the proteasome (mild stress) in ISCs which resulted in cell cycle arrest because of the presence of protein aggregates. Remarkably, when the ISCs cleared the aggregates they were able to resume cell cycle and thus overcome this major hurdle (Rodriguez-Fernandez et al., 2019).

The human chaperome is formed by 332 chaperones and co-chaperones and each family of chaperones is essential for cell viability indicating that they have non-overlapping functions (Brehme et al., 2014). Indeed some planarian chaperones and also components of the ubiquitin-proteasome system have been linked to stem cell regulation (*prohibitin-1* and *-2*, *mortalin*, *hsp40* and *hsp60*) (Conte et al., 2009; Fernandez-Taboada et al., 2011; Henderson et al., 2015; Isolani et al., 2012; Rossi et al., 2014; Strand et al., 2018; Wang et al., 2019) and our data validates their enrichment in stem cells. Although we currently do not know whether these effects are nutrient dependent, cluster 1 data set predicts that at least *prohibitin-2* (Rossi et al., 2014) and one *hsp40* (Fernandez-Taboada et al., 2011) could also have a nutrient-dependent effect on stem cells. It would be interesting to know whether there are other chaperones apart from TRiC which are specifically required in stem cells during starvation and to determine the mechanism through which stem cells are protected under these conditions. Our work establishes planarians as an excellent *in vivo* model to investigate functions of proteins in metabolism.

We have investigated the role of TRiC in starved stem cells. Mechanistically, we have found that *cct3A*, a subunit of TRiC is necessary to allow mitosis during the response of stem cells to amputation. *cct3A* RNAi leads to gross nuclear alterations, which include asymmetrical karyokinesis, abnormal organization of the spindle poles and chromosome lagging and an accumulation of stem cells in anaphase. All this leads to the cell death observed at wound sites at 4dR. These results highlight the importance of proteostasis to maintain genome integrity. Although it is known that altered proteostasis and genome instability are linked to an increased risk of cancer (Adams et al., 2015; Dai and Sampson, 2016), it is not known whether proteostasis is essential for protecting the genome. There are only few evidences indicating a possible role of proteostasis in regulating genome stability. For instance, it has been reported that ER stress and thus activation of UPR^ER^ induced by tunicamycin or glucose deprivation, suppresses DNA double-strand break repair in cancer cells by stimulating the degradation of Rad51 (Yamamori et al., 2013). Also Hsp70 has been linked to genome stability in mouse embryonic fibroblasts under heat shock stress (Hunt et al., 2004) and Hsp110 is associated to genome instability in cancer cells (Dorard et al., 2011). Interestingly recent studies in planarians and *Drosophila* have shown that the heat shock protein DNAJA1 and HSP90 interact with PIWI proteins suggesting a possible role of these chaperones in suppressing transposition (Gangaraju et al., 2011; Wang et al., 2019). Although TRiC has a potential role in cancer development by modulating the folding of client proteins related to oncogenesis, cell cycle, and cytoskeleton (i.e. actin and tubulin) (reviewed in (Roh et al., 2015), a direct role in genome stability of stem cells is still missing. Remarkably, double RNAi for two of the main transcription factors of UPR^ER^ *xbp1* and *atf6* phenocopies *cct3A* RNAi hinting towards a previously unknown crosstalk between the cytosol and the ER during starvation in stem cells in the context of regeneration. Thus it is unlikely that the mechanisms through which they regulate genome integrity are related to specific clients that depend on the folding by TRiC. Nevertheless, this altogether indicates that the main outcome after *ccts* or UPR^ER^ down-regulation during starvation is related to genome integrity in stem cells. Moreover we show that starvation protects the planarian stem cell genome and thus possibly contributes to the capacity of planarians to circumvent cancer. These results also suggests that the protective effect that fasting confers to stem cells upon chemotherapy (Nencioni et al., 2018) could be related to increased proteostasis.

It is known that LDs accumulate under various stress conditions. For instance, mice injected with the ER-stress inducer tunicamycin develop hepatic steatosis which is even more severe and showing LDs accumulation when any of the UPR branches is knocked-down (Rutkowski et al., 2008; Yamamoto et al., 2010). Although the current view is that LDs play a protective role storing toxic lipids to avoid their accumulation within the ER membrane (Hapala et al., 2011), our results support an alternative view in which stress would activate the UPR and then lipid catabolism to provide stem cells with the required energy to progress with the cell cycle. Indeed prior work has shown that fasting enhances intestinal stem cell function through activation of fatty acid oxidation (Mihaylova et al., 2018). Several of our results imply a role of TRiC and UPR activation in this process. One single feeding during the fasting period is sufficient to prevent *cct3A(RNAi)* and *xbp1/atf6(RNAi)* phenotypes and allows normal mitosis and regeneration thus indicating that energy production and/or the supply of nutritional building blocks for cellular anabolism prevent the phenotype. Furthermore, the RNA-seq data indicates that both *cct3A* RNAi and *xbp1/atf6* RNAi regulate lipid metabolism. Indeed for *cct3A* RNAi we observed less LDs in intestinal branches at 4dR. The RNA-seq data indicates that *cct3* RNAi up-regulates LD biogenesis, fatty acid β-oxidation and sphingolipids biosynthesis while lowering ATP levels in the planarians, which overall suggest that starved *cct3A(RNAi)* planarians cannot generate sufficient energy and possibly fail to provide sufficient fatty acids and/or cholesterol for membrane expansion in order to pass mitosis, a highly energy demanding process (Kalucka et al., 2015). In *xbp1/atf6(RNAi)* planarians, RNA-seq data indicates that it down-regulates LD biogenesis, LD breakdown and fatty acid β-oxidation which correlates with the observed accumulation of LDs in differentiated and stem cells of the blastemas and suggest that *xbp1/atf6(RNAi)* animals cannot mobilize LDs to obtain energy. *cct3A* RNAi in contrast *to xbp1/atf6* RNAi does not show ATGL or other lipases differentially down-regulated. However we found an homolog for *TFEB*, which was shown to promote lipophagy and regulate the lysosome compartment (Settembre et al., 2013), to be down-regulated in starved *cct3A(RNAi)* animals. Remarkably *ccts* are also known to be required for autophagosome degradation through regulating lysosomal acidification in HeLa cells and mouse primary cortical neurons (Pavel et al., 2016). It is known that autophagy is required for LD breakdown (Singh et al., 2009). Indeed the current view on the mechanism of LD breakdown is that ATGL and possibly other lipases are upstream regulators of lipophagy and that lipophagy (rather than the lipases) is responsible for the bulk breakdown of LDs (Schulze et al., 2017). Altogether, our findings suggest that *cct3A* RNAi down-regulates LD degradation by targeting the lysosomal compartment and thus interfering with the breakdown of LDs in the autophagosome, whereas *xbp1/atf6* RNAi does the same by targeting the lipases related to LD breakdown. Therefore *cct3A* may impair a double regulation on LD breakdown dependent and independent of the UPR which may explain why we detect less ATP in *cct3(RNAi)* animals, and also the accumulation of pinocytic vesicles, which has been often seen as a last resource to obtain energy from the extracellular space, at least in cancer cells (Commisso et al., 2013).

We have previously shown that starvation is able to rejuvenate the stem cell pool in terms of telomere length through downregulation of mTOR signalling (Iglesias et al., 2019). This work indicates that starvation also regulates stem cells by increasing their stress and metabolic resistance through proteostasis which protects and allows the regenerative mitotic responses during starvation. Our data contributes to the understanding of stem cell biology, proteostasis and ageing and consolidates the planarian as a feasible model system which to study the consequences of fasting in stem cells.

## Experimental Procedures

### Planarian husbandry

Planarians used in this work belong to the species *S. mediterranea* asexual biotype. All animals were maintained at 19°C in 1x Montjuïc Salts (1.6mM NaCl, 1mM CaCl2, 1mM MgSO4, 0.1mM MgCl2, 0.1mM KCl, 0.1gr NaHCO3) and fed with organic veal liver.

### Flurorescence-activated cell sorting (FACS)

Planarian dissociation and cell population analysis were performed as described previously (Hayashi et al., 2006). Details can be found at Supplemental Experimental Procedures.

### RNA-seq

Library preparation was done using the Illumina kit TruSeq. Sequencing was done on an Illumina HiSeq2500 in 51 cycle, single-end, high-throughput mode at the Core facility DNA sequencing at FLI. Details can be found in Supplemental Experimental Procedures.

### RNAi experiments

Templates with T7 promoters appended to both strands were generated. Double-stranded RNA (dsRNA) was synthesized by *in vitro* transcription following MEGAscript RNAi kit (Ambion) instructions. dsRNA was injected into the planarian as previously described (Gonzalez-Estevez et al., 2012b). Control animals were injected with *gfp* dsRNA, a sequence not present in the planarian genome. The different protocols used are specified in the “Results” section. More details can be found in Supplemental Experimental Procedures.

### Whole-mount *in situ* hybridization

Whole-mount *in situ* hybridization was carried out as described previously (King and Newmark, 2013) using an InsituPro VSi (Intavis). Hapten-labelled RNA probes were prepared by using an *in vitro* RNA labelling kit (Roche).

### Statistical analysis

In all the manuscript error bars are s.d. from the mean and asterisks indicate P < 0.05 (one asterisk), P < 0.01 (two asterisks) or P < 0.001 (three asterisks) and n.s. indicates not significant using two-tailed Student’s test with equal variance. The number of replicates is indicated in figures and figure legends. Wald-test was used to identify differentially expressed genes (DEGs). Significance was determined by q-value (false discovery rate (FDR)) < 0.1 for pairwise comparisons for the different time points of starvation (7dS vs 1dS, 30dS vs 1dS and 30dS vs 7dS) in the X1. Significance was determined by q-value < 0.01 for pairwise comparisons X1 vs Xins at 1dS, 7dS and 30dS. Significance was determined by q-value < 0.05 in the *cct3(RNAi)* and *xbp1/atf6(RNAi)* RNA-seq analysis. More details on the RNA-seq statistics can be found in Supplemental Experimental Procedures.

### Data availability

Sequences of *cct6* and *cct8A* have been previously deposited in GenBank: MF669568 and MF669570, respectively (Counts et al., 2017). Sequences of *xbp1*, *atf6*, *cct3A*, *cct1A*, *cct2*, *cct4B*, *cct5*, *cct7* have been deposited in GenBank with the accession numbers MN171093-MN171100. RNA-Seq data has been deposited in GEO with the accession numbers GSE134148, GSE134105 and GSE134013.

## More methods can be found in Supplemental Experimental Procedures

## Acknowledgements

We would like to thank the Histology, Imaging, DNA Sequencing and Flow Cytometry Core Facilities from the Leibniz Institute on Aging-Fritz Lipmann Institute (FLI) for their technical support. We are grateful to J. Solana, M. Rodriguez-Orejuela and all past and current members in CGE lab for discussions, suggestions and/or help on the project. We would like to thank M. Iglesias for critical reading of the manuscript and technical support. We are grateful to T. Adell for the protocol for staining with anti-α-tubulin and to Prof. K. Watanabe and H. Orii for providing the VC1 antibody. CGE was funded by the Leibniz Institute on Aging-Fritz Lipmann Institute (FLI). The FLI is a member of the Leibniz Association and is financially supported by the Federal Government of Germany and the State of Thuringia. OGG was funded by a LGSA (Leibniz Graduate School on Ageing and Age Related Diseases) scholarship.

## Author contributions

OGG, DAF and CGE performed most of the experiments with the help of AT; OGG performed computational analysis; AK and KLR helped in the project; SP performed the clustering analysis; AS performed the electron microscopy experiments; OGG,

DAF and CGE designed experiments and analyzed the data; CGE directed the project; CGE wrote the manuscript with help of OGG, DAF, AS, AK and KLR.

