## Supplemental Information for "TRiC activates the unfolded protein response and protects starved stem cells by modulating energy and lipid metabolism during planarian regeneration"

**This PDF file includes:**

Supplemental Experimental Procedures
Figure S1 to S4
Captions for Tables S1 to S5 (provided as separate Excel files)

**Other Supplemental Materials for this manuscript include:**
Tables S1 to S5

### Supplemental Experimental Procedures

#### Starvation and feeding experiments

In experiments involving feeding, all animals were observed to make sure they ate. Food was given in excess and removed after 2 hours. For RNA-seq experiments planarians at 1dS, 7dS and 30dS had the same area 4 $mm^2$ (5-5.5 mm length at 7dS and 1dS and 5.5- 6mm length at 30dS). Graph paper placed under the Petri dish was used to pre-select animals and the final selection was done after measuring the areas with the Leica Application Suite (Leica) on photographs of live planarians taken under a stereomicrospe coupled with a Leica camera MC170 HD (Leica). For RNAi experiments planarians around 5 mm were selected by use of graph paper.

#### Flurorescence-activated cell sorting (FACS)

Planarians were cut into small pieces on ice and cell dissociated in the presence of papain (Merck; final concentration 1mg/ml) during 15 min at room temperature (Moritz et al., 2012). Cells were then filtered through a 40$\mu$m filter, counted and resuspended in staining solution containing the cytoplasmic dye Calcein-AM (Biotium; final concentration of 0.5 $\mu$g/ml) and the nuclear dye Hoechst 33342 (Thermo Fisher Scientific; final concentration 30$\mu$g/ml) in order to isolate X1, X2 and Xins populations. Staining was performed in the dark at 25ºC, with continuous shaking. Propidium Iodide (final concentration 1$\mu$g/ml) was added 1 minute before flow cytometry analysis to discard dead cells. Around 250,000 events were sorted per sample using BD FACSAria III or BD FACS Fusion. Cells were put into tubes containing Trizol LS (Ambion) and RNA was extracted following manufacturer's instructions.

#### RNA-seq

4 replicates per FACS population (X1, X2 and Xins) and per time point (1dS, 7dS and 30dS) were sequenced: 36 samples (2,304 million of non-ribosomal reads). 3 replicates per RNAi condition (*cct3A(RNAi)*, *gfp(RNAi)*, *xbp1/atf6(RNAi)* and *gfp/gfp(RNAi)* were sequenced: 12 samples (516 million of non-ribosomal reads). For transcriptomic analysis, the Dresden *Schmidtea mediterranea* transcriptome (version 4) was used as reference (PlanMine) (Rozanski et al., 2019). Kallisto (version 0.43.0) (Bray et al., 2016) was used to perform read pseudo-alignment and quantification with the parameters –l 200 –s 20 –b 100. Previously, only reads mapping to the planarian reference transcriptome were extracted by using Kallisto and the parameter -F 4. The portion of planarian ribosomal RNA contamination was identified by mapping all reads with Bowtie2 (Langmead and Salzberg, 2012) against a pool of platyhelminthes rRNA index. IDs numbers from Dresden transcriptome of version 4 are equivalent to version 6 (Planmine). For example, dresden_comp15_c0_seq1 is equivalent to dd_Smed_v6_15_0_1. Differential expression analysis was performed with the R package Sleuth (version 0.28.1) (Pimentel et al., 2017) with the filtered reads mapping to the reference transcriptome. Wald-test was used to identify differentially expressed genes (DEGs). Significance was determined by q-value (false discovery rate (FDR) < 0.1 for pairwise comparisons for the different time points of starvation (7dS vs 1dS, 30dS vs 1dS and 30dS vs 7dS) in the X1. For pairwise comparisons X1 vs Xins at 1dS, 7dS and 30dS all TPMs were normalized including also X2 values and the significance was determined by q-value < 0.01. Significance was determined by q-value < 0.05 in the *cct3(RNAi)* and *xbp1/atf6(RNAi)* RNA-seq analysis. Downstream analysis was done with the R software. Heatmaps were done with the R package "Pheatmap". Clustering was performed using the software package Mfuzz (Kumar and M, 2007). Genes with a value lower than 0.35 for cluster assignment are not displayed in the plots. Gene ontology enrichment was done creating lists of DEGs in PlanMine

by using default parameters (test Benjamini Hochberg; p-value < 0.05). Redundant GO terms were removed from the list by using REVIGO with small similarity parameter (Supek et al., 2011).

**RNAi experiments**

The following primers were used to generate the templates for dsRNA production:

Smed-cct1A-F: 5´-ACCTGGCTATTGGTGGAGAAAG-3´

Smed-cct1A-R: 5´-GGCATTTGTTGGGAAGCTATTG-3´

Smed-cct2-F: 5´-TGACAACCCTGCTGCTAAAATC-3´

Smed-cct2-R: 5´-TGGCAATCTTTTCGACCTTTTC-3´

Smed-cct3A-F: 5´-CGTCGTTTTGAGTGGAGTTTTG-3´

Smed-cct3A-R: 5´-TTGATATTGCCATCTCCAATGC-3´

Smed-cct4B-F: 5´-CTCCAATAGCAGTTGATGCAG-3´

Smed-cct4B-R: 5´-GGCCAGGATTAACAATTCCACT-3´

Smed-cct5-F: 5´-TCACTCGGACCAAAGGGATTAG-3´

Smed-cct5-R: 5´-TGGAGGTTCAAAAGCACAAGTC-3´

Smed-cct6-F: 5´-GAGTACGCAAAGTCGCAAAATG-3´

Smed-cct6-R: 5´-TTTCGGCATATCTGGATGTCTG-3´

Smed-cct7-F: 5´-AAATGTGCTTCCACTGCTCTCA-3´

Smed-cct7-R: 5´-ACCGCTCTGTTTCCTCCATAAA-3´

Smed-cct8-F: 5´-AATGGCTGCTCAACAACAAGAG-3´

Smed-cct8-R: 5´-TGCGTTTCTTCTCCACGACTAA-3´

Smed-xbp1-F: 5´-TAGGTGGGAATGGTATGGGAAA-3´

Smed-xbp1-R: 5´-CACAACCAAACTCTGACATTTCG-3´

Smed-atf6-F: 5´-AAGCCAGTTGTTAAGCCAGAAA-3´

Smed-atf6-R: 5´-CCATGATAACCGGGAAATGAAGA-3´

**Immunohistochemistry**

Whole-mount immunohistochemistry was performed as previously described (Cebria and Newmark, 2005). The following primary antibodies were used: anti-VC-1 (Sakai et al., 2000) (diluted 1/15000; kindly provided by Professor K. Watanabe and H. Orii); anti-TMUS13 (Cebria et al., 1997) (diluted 1/20; kindly provided by Professor R. Romero); anti-acetylated-tubulin (Reddien et al., 2007; Robb and Sanchez Alvarado, 2002) (diluted 1:200, Sigma; clone 6-11B-1); and anti-Histone H3 phosphorylated at serine 10 (H3P, diluted 1/500, Santa Cruz; sc-8656-R). Nuclei were stained with DAPI (1ug/ml). Double immunohistochemistry on dissociated cells was performed as previously described (de Sousa et al., 2018). An anti-Histone H3 phosphorylated at serine 10 (H3P, diluted 1/500, Santa Cruz; sc-8656-R) and anti-Tubulin Tyrosin (diluted 1/400; Sigma; T9028) were used.

**Real Time PCR (qPCR)**

RNA was extracted using Trizol (Ambion) for whole planarians and Trizol LS (Ambion) for FACs sorted cells. cDNA was obtained from 200 ng (from sorted cells) or 1 $\mu$g (from whole animals) of total RNA by using MMLV Reverse Transcriptase (Promega). As internal control

Elongation Factor 2 (EF2) and the transcript with ID 5685 from Dresden transcriptome (Planmine) were used. Each qPCR was performed with three biological replicates. Five animals were used per replicate, and each sample was replicated three times in each real-time PCR experiment. PCR reactions were performed using the iTaq Universal SYBER® Green Supermix (BIO-RAD). Reactions were aliquoted using a QiAgility robot (Qiagen) and analyzed with a 7500 Real Time PCR System (Applied Biosystems). The following gene specific oligos were designed from non-overlapping regions of the gene that were non-overlapping with sequence used for dsRNA:

qWi1PPF: 5'-GCAGAGAAACGGAAGTAATAGAG-3'

qWi1PPR: 5'-ATCCAATCCTACAATCATAGTCGG-3'

xbp1-QF: 5´-ATTGGAAACTCCCATTTGTGAC-3´

xbp1-QR: 5´-AGAATTATCTGGCTGACTTTGG -3´

atf6-QF: 5´- TAAGAAATCGAAATTCCGGGAC-3´

atf6-QR: 5´- AAGCGTATAATCCTTGGTTCTG-3´

cct3A_QF: 5´- GCAATATCAACTCACTTGACCCT-3´

cct3A_QR: 5´- TAACCTTAACACCTTTGCTCCA-3´

cct4B_QF: 5´- CAGAGTTAAGAAATAGACATGCCAG-3´

cct4B_QF: 5´- ACTGTGATTGATAAGGAGTGGT-3´

cct7_QF: 5´- AATTCCACGACAATTATGCGAG-3´

cct7_QR: 5´- CTTCATTAAGAATGTCGACTCCAC-3´

**Transmission electron microscope analysis**

Planarians were fixed in 4% paraformaldehyde and 2.5% glutaraldehyde solution in 0.1M cacodylate buffer, pH 7.2, for 3h at room temperature. Planarians were post-fixed in 1% osmium tetroxide in 0.1M cacodylate buffer for 2h at room temperature. After rapid dehydration in a graded series of ethanol and propylene oxide, samples were embedded in an "Epon-Araldite" mixture. Ultrathin sections, obtained by a diamond knife on an Ultracut Reichert-Jung ultramicrotome, were placed on Formvar-carbon coated nickel grids, stained with uranyl acetate and lead citrate and observed with a Jeol 100 SX transmission electron microscope. The quantification was performed by counting the number of differentiated and neoblast-like cells with lipids or vacuoles per blastema.

**Measurement of ATP levels**

ATP levels were measured by using ATP bioluminescence assay kit HS II (Roche) according to the manufacturer's instructions. Regenerating trunks were incubated in cell lysis reagent and boiled for 15 minutes at 100ºC. Then, samples were sonicated for 4 cycles of 30''. Luciferase reagent was added to the samples and luminescence was measured by the Mithras LB940 plate reader (Berthold Technologies) using the injection function. Three biological replicates consisting of five trunks at 48hR were used per condition, and each sample was replicated three times in each ATP experiment.

**Whole-mount TUNEL**

Whole-mount TUNEL was performed as previously described (Stubenhaus and Pellettieri, 2018).

**Oil Red O staining on cryosections**

5 planarians per condition were killed in 2% HCl in Holtfreter 5/8, washed 2x 5min in Holtfreter 5/8 and fixed O/N in 4% PFA in PBST (0.1% Triton X-100) at 4°C. After 2 washes in PBST and 2 washes in PBS of 5 min each they were incubated in 15% sucrose in PBS for 20 min and in 30% sucrose in PBS O/N at 4°C before being embedded in OCT and cryosectioned for 14 µm thickness. After 3 x 5 min washes in PBS the sections were first immersed in 60% isopropanol for 2 min and then for 20 min in Oil Red O solution in isopropanol while shaking. Oil Red O in isopropanol is prepared as follows: a 0.5g/100mL isopropanol stock solution equilibrated for several days was freshly diluted to 60% with water and rocked for at least 1h then vacuum-filtered. Unspecific precipitate was removed by quickly dipping into 60% isopropanol. After a 5 min wash in PBS the nuclei were counterstained using 5ug/ml DAPI in PBS for 5 min. 70% glycerol in PBS was used as mounting medium.

**Imaging and quantifications**

Z-stacks were acquired with a Zeiss ApoTome.2 equipped with a Zeiss Axiocam 503 mono (Carl Zeiss, Jena) (immunohistochemistry on dissociated cells), a Zeiss LSM 710 ConfoCor 3 microscope (Carl Zeiss, Jena) (whole-mount immunohistochemistry, Oil Red O staining on cryosections and whole-mount TUNEL) or a Zeiss AXIO Zoom.V16 (Apotome.2) equipped with Axiocam 506 mono and color (Carl Zeiss, Jena) (H3P whole-mount immunohistochemistry and whole-mount FISH). Images were processed using Fiji (Schindelin et al., 2012) and Adobe Photoshop CS6/Adobe CC software. H3P quantifications were done on whole planarians using Object Counter 3D plugin from Fiji (Schindelin et al., 2012).

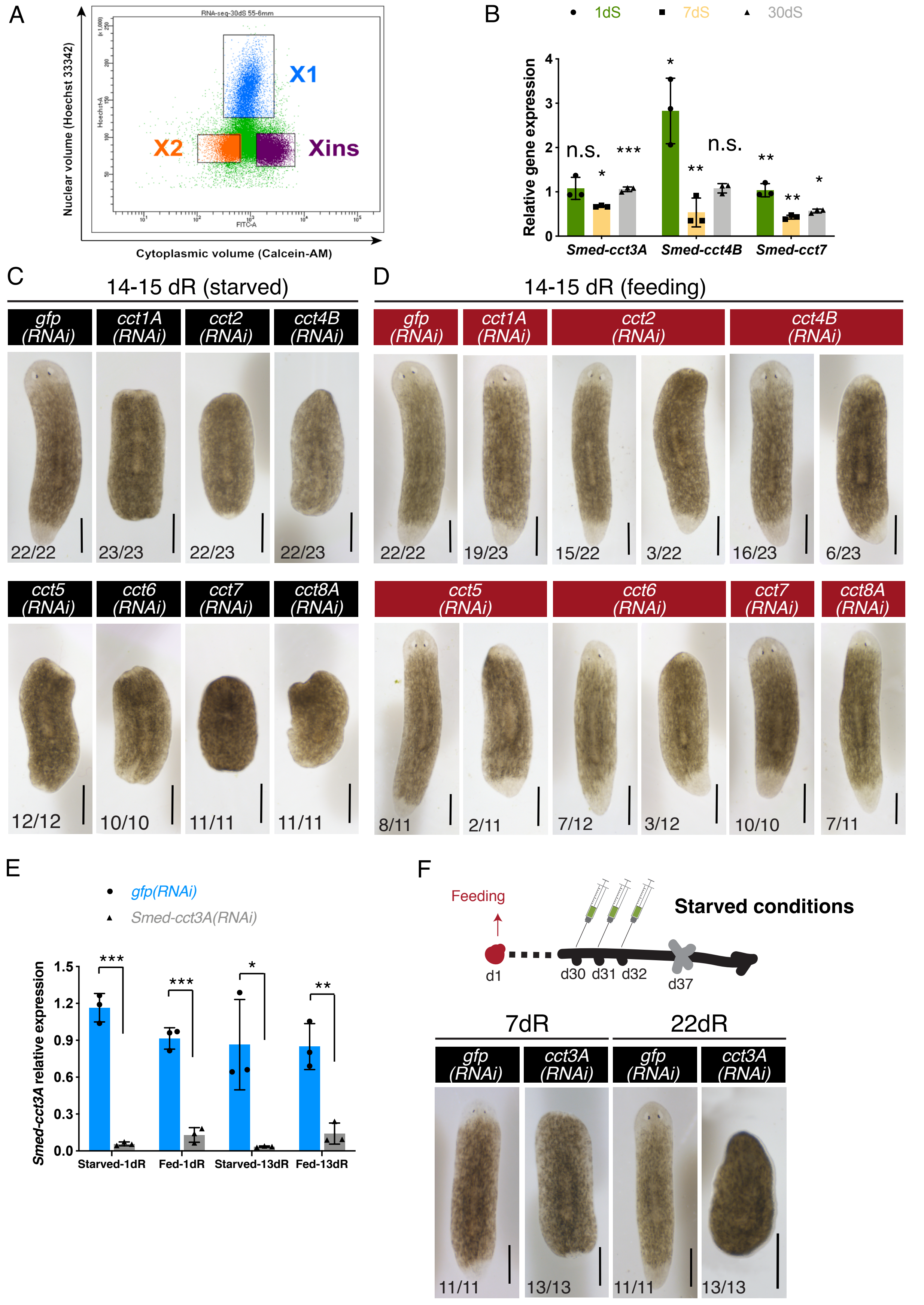
A
Nuclear volume (Hoechst 33342)
Cytoplasmic volume (Calcein-AM)
RNA-seq-30dS 55-6mm
X1
X2
Xins
B
1dS
7dS
30dS
Relative gene expression
n.s.
*
***
**
Smed-cct3A
Smed-cct4B
Smed-cct7
C
14-15 dR (starved)
gfp (RNAi)
cct1A (RNAi)
cct2 (RNAi)
cct4B (RNAi)
22/22
23/23
22/23
22/23
cct5 (RNAi)
cct6 (RNAi)
cct7 (RNAi)
cct8A (RNAi)
12/12
10/10
11/11
11/11
D
14-15 dR (feeding)
gfp (RNAi)
cct1A (RNAi)
cct2 (RNAi)
cct4B (RNAi)
22/22
19/23
15/22
3/22
16/23
6/23
cct5 (RNAi)
cct6 (RNAi)
cct7 (RNAi)
cct8A (RNAi)
8/11
2/11
7/12
3/12
10/10
7/11
E
gfp(RNAi)
Smed-cct3A(RNAi)
Smed-cct3A relative expression
Starved-1dR
Fed-1dR
Starved-13dR
Fed-13dR
F
Feeding
Starved conditions
d1
d30
d31
d32
d37
7dR
22dR
gfp (RNAi)
cct3A (RNAi)
gfp (RNAi)
cct3A (RNAi)
11/11
13/13
11/11
13/13

**Figure S1. TRiC is necessary for blastema formation specifically in starved planarians.** (A) FACS profile of planarian cells according to nuclear and cytoplasmic volume. The RNA-seq compared X1 cells (dividing stem cells) from planarians at different nutritional status. (B) Relative expression of *cct* transcripts at 1, 7 and 30 days of starvation in X1 (stem cells). Error bars are s.d. from the mean. Asterisks refer to the condition just before and 1dS refer to 30dS and indicate P < 0.05 (one asterisk), P < 0.01 (two asterisks), P < 0.001 (three asterisks) and n.s. indicates not significant using two-tailed Student's test with equal sample variance. n = 3 replicates (5 planarians each) per time point. (C) RNAi injections schedule in starved conditions (14dS when the amputation is performed). Live images show that *RNAi* for any *cct* gene leads to minimal blastema formation compared to controls at the time points shown. At the bottom are the number of planarians with the phenotype shown. The remaining planarians in *cct2(RNAi)* are dead and like controls in *cct4B(RNAi)*. (D) RNAi injections schedule in feeding conditions (one extra feeding respect to C is introduced one day prior to injections). Live images show that RNAi for any *cct* gene leads to most planarians regenerating as controls. At the bottom are the number of planarians with the phenotype shown. The remaining planarians are dead by the time point of regeneration shown. (E) Relative expression of *cct3A* in *cct3A(RNAi)* and control planarians injected either under starved or feeding conditions at 1 day and 13 days of regeneration. Error bars are s.d. from the mean. Asterisks indicate P < 0.05 (one asterisk), P < 0.01 (two asterisks), P < 0.001 (three asterisks) and n.s. indicates not significant using two-tailed Student's test with equal sample variance. n = 3 replicates (5 planarians each) per time point. (F) RNAi injections schedule in starved conditions. Planarians are at 37 days of starvation when the amputation is performed (indicated by the grey cross at day 37). Live images show that *cct3A(RNAi)* planarians form a minimal blastema when compared to controls at the time points shown. At the bottom are the number of planarians with the phenotype shown. dR, days of regeneration; dS, days of starvation. Scales, 500 μm.

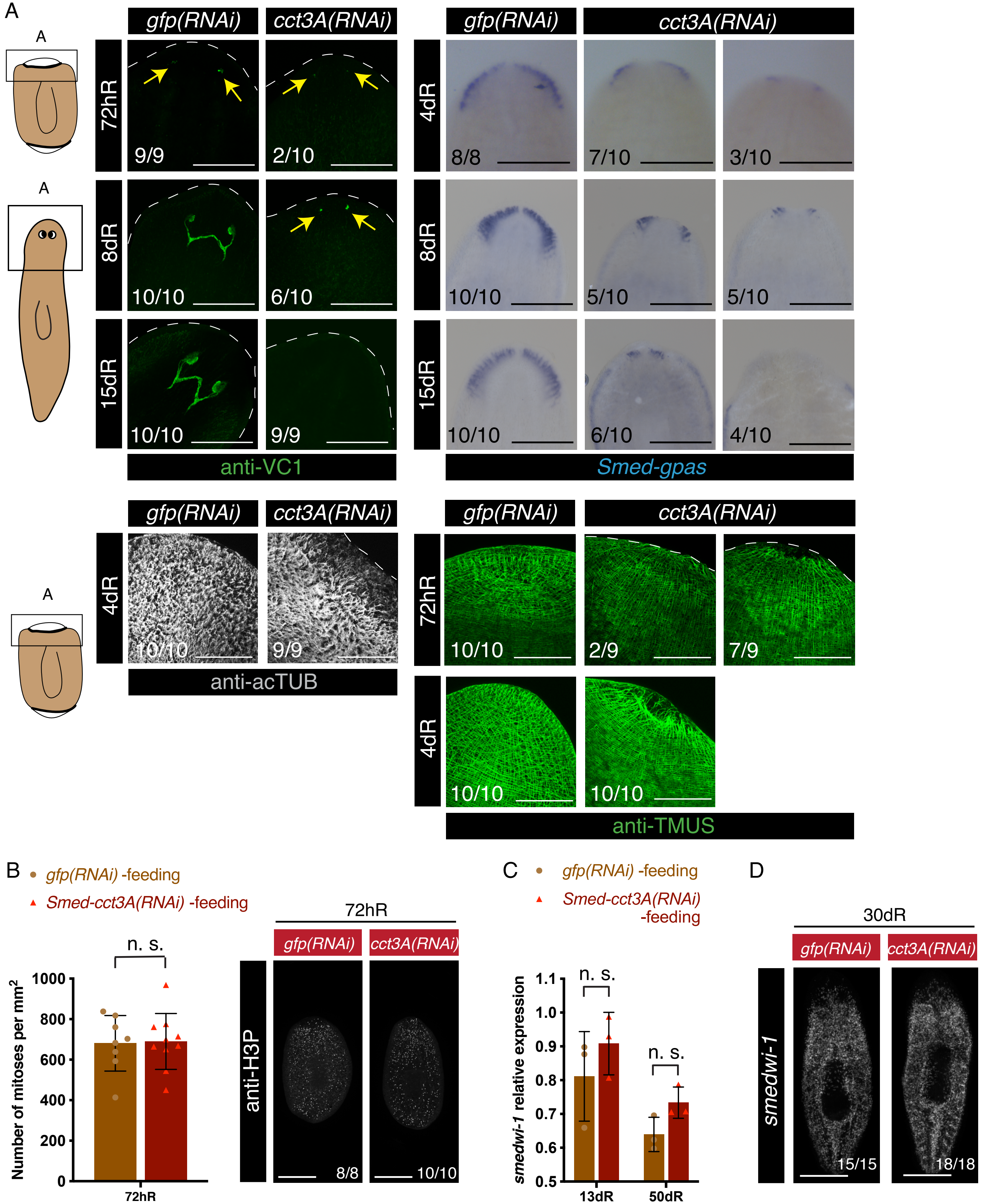

A
gfp(RNAi)
cct3A(RNAi)
72hR
8dR
15dR
4dR
9/9
2/10
10/10
6/10
10/10
9/9
anti-VC1
8/8
7/10
3/10
10/10
5/10
5/10
10/10
6/10
4/10
Smed-gpas
10/10
9/9
anti-acTUB
10/10
2/9
7/9
10/10
10/10
anti-TMUS
B
gfp(RNAi) -feeding
Smed-cct3A(RNAi) -feeding
n. s.
Number of mitoses per mm$^2$
1000
800
600
400
200
0
72hR
anti-H3P
8/8
10/10
C
smedwi-1 relative expression
1.1
1.0
0.9
0.8
0.7
0.6
0.5
13dR
50dR
D
30dR
smedwi-1
15/15
18/18

**Figure S2. *cct3A* RNAi leads to minimal differentiation during regeneration in starved conditions while in feeding conditions mitoses and stem cells are as in controls.** (A) The squares at the cartoons indicate the regions shown in the panels. The images show minimal differentiation of eyes (anti-VC1), brain (*Smed-gpas*), epidermal cilia (anti-acTUB) and muscle (anti-TMUS) in anterior wounds of *cct3A* RNAi compared to controls. At the bottom are the number of planarians with the phenotype shown. Arrows indicate the differentiating eyes. (B) The graph shows the mitotic numbers during 72 hours of regeneration in feeding conditions for *cct3A* RNAi and controls. Error bars are s.d. from the mean and n.s. indicates not significant; n $\geq 8$ planarian per time point. Maximum projections of representative trunks labelled with anti-H3P at 72 hours of regeneration. On the bottom, the number of planarians with the phenotype shown from the total is displayed. (C) Relative expression of *smedwi-1* at 13 and 50 days of regeneration after either *cct3A* or *gfp* RNAi under feeding conditions. Error bars are s.d. from the mean and n.s. indicates not significant using two-tailed Student's test with equal sample variance. n = 3 replicates (5 planarians each) per time point. (D) Maximum projections of representative trunks after FISH for *smedwi-1* at 30 days of regeneration in *cct3A* RNAi and controls under feeding conditions show that the expression levels are similar. On the bottom, the number of planarians with the phenotype shown from the total is displayed. hR, hours of regeneration; dR, days of regeneration. Scales, 300 µm (VC1 images), 500 µm (*gpas* images, B and D), 150 µm (TMUS and AC-TUB images).

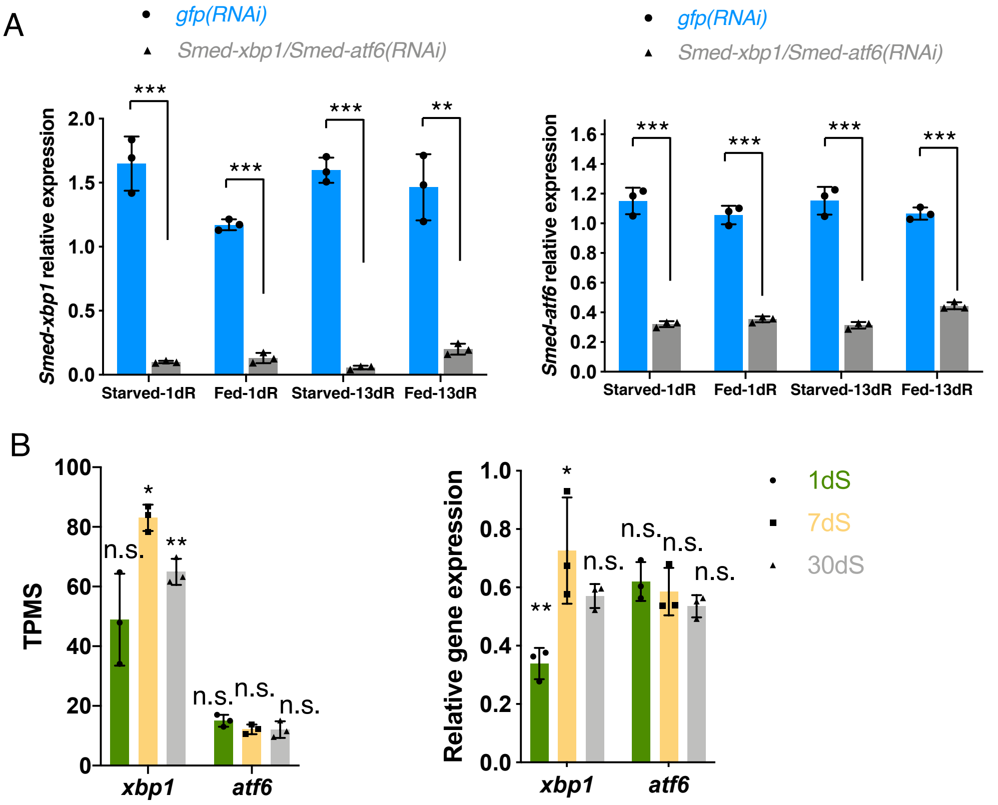
A
gfp(RNAi)
Smed-xbp1/Smed-atf6(RNAi)
Smed-xbp1 relative expression
2.0
1.5
1.0
0.5
0.0
***
***
***
**
Starved-1dR
Fed-1dR
Starved-13dR
Fed-13dR
gfp(RNAi)
Smed-xbp1/Smed-atf6(RNAi)
Smed-atf6 relative expression
1.6
1.4
1.2
1.0
0.8
0.6
0.4
0.2
0.0
***
***
***
***
Starved-1dR
Fed-1dR
Starved-13dR
Fed-13dR
B
TPMS
100
80
60
40
20
0
n.s.
*
**
n.s.
n.s.
n.s.
xbp1
atf6
Relative gene expression
1.0
0.8
0.6
0.4
0.2
0.0
**
*
n.s.
n.s.
n.s.
n.s.
xbp1
atf6
1dS
7dS
30dS

**Figure S3. Expression of *xbp1* and *atf6* after *xbp1/atf6* RNAi and at different nutritional status**. (A) Relative expression of *xbp1 and atf6* at 1 day and 13 days of regeneration after *xbp1/atf6(RNAi)* and *gfp*(RNAi) during either starving or feeding conditions. The graphs show that *xbp1/atf6* RNAi down-regulates *xbp1* and *atf6* during starvation and feeding at both time points. Error bars are s.d. from the mean. Asterisks indicate P < 0.01 (two asterisks), P < 0.001 (three asterisks) and n.s. indicates not significant using two-tailed Student's test with equal sample variance. n = 3 replicates (5 planarians each) per time point. (B) Expression levels of *xbp1 and atf6* at 1 day, 7 days and 30 days of starvation in TPMs (transcripts per million) and relative expression of *xbp1 and atf6* at 1 day, 7 days and 30 days of starvation. Error bars are s.d. from the mean. Asterisks refer to the condition before and 1dS refers to 30dS and indicate P < 0.05 (one asterisk), P < 0.01 (two asterisks) and n.s. indicates not significant using two-tailed Student's test with equal sample variance. n = 3 replicates (5 planarians each) per time point.

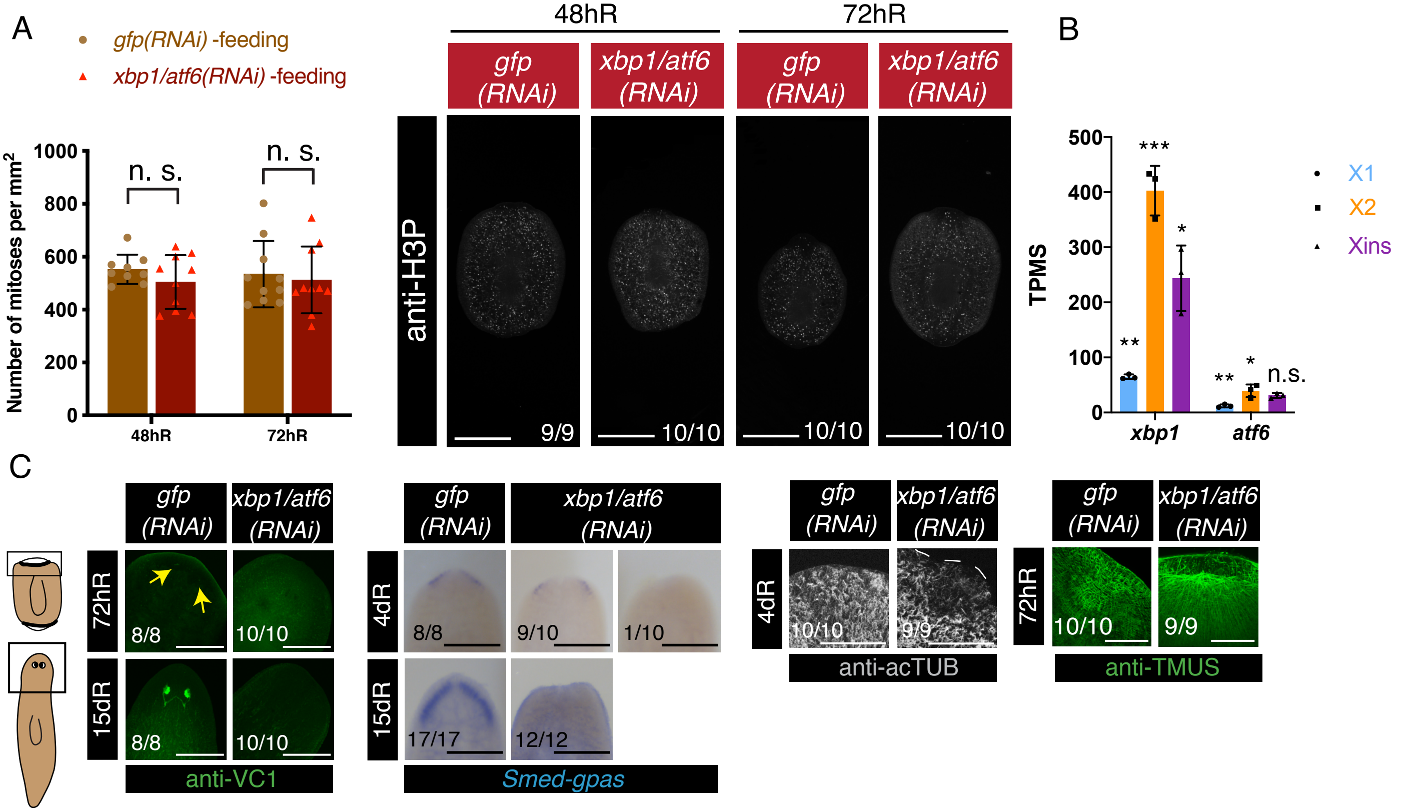
A
gfp(RNAi) -feeding
xbp1/atf6(RNAi) -feeding
Number of mitoses per mm$^2$
1000
800
600
400
200
0
n. s.
n. s.
48hR
72hR
48hR
72hR
gfp (RNAi)
xbp1/atf6 (RNAi)
gfp (RNAi)
xbp1/atf6 (RNAi)
anti-H3P
9/9
10/10
10/10
10/10
B
TPMS
500
400
300
200
100
0
***
*
**
**
*
n.s.
X1
X2
Xins
xbp1
atf6
C
gfp (RNAi)
xbp1/atf6 (RNAi)
72hR
15dR
8/8
10/10
8/8
10/10
anti-VC1
gfp (RNAi)
xbp1/atf6 (RNAi)
4dR
15dR
8/8
9/10
1/10
17/17
12/12
Smed-gpas
gfp (RNAi)
xbp1/atf6 (RNAi)
4dR
10/10
9/9
anti-acTUB
gfp (RNAi)
xbp1/atf6 (RNAi)
72hR
10/10
9/9
anti-TMUS

**Figure S4.** ***xbp1/atf6*** **RNAi leads to minimal differentiation during regeneration in starved conditions while in feeding conditions mitoses are as in controls.** (A) The graph shows the mitotic numbers during 48 hours and 72 hours of regeneration in feeding conditions for *xbp1/atf6* RNAi and controls. Error bars are s.d. from the mean and n.s. indicates not significant using two-tailed Student's test with equal sample variance; $n \geq 9$ planarians per time point. Maximum projections of representative trunks labelled with anti-H3P at 48 hours and 72 hours of regeneration. On the bottom, the number of planarians with the phenotype shown from the total is displayed. (B) Expression levels of *xbp1 and atf6* at 30 days of starvation in the X1, X2 and Xins FACS populations in TPMs (transcripts per million). Error bars are s.d. from the mean. Asterisks refer to the condition before and X1 refers to Xins and indicate $P < 0.05$ (one asterisk), $P < 0.01$ (two asterisks), $P < 0.001$ (three asterisks) and n.s. indicates not significant using two-tailed Student's test with equal sample variance. (C) The squares at the cartoons indicate the regions shown in the panels. The images show no differentiation of eyes (anti-VC1), and minimal differentiation of brain (*Smed-gpas*), epidermal cilia (anti-acTUB) and muscle (anti-TMUS) in anterior wounds of *xbp1/atf6* RNAi compared to controls. At the bottom are the number of planarians with the phenotype shown. Arrows indicate the differentiating eyes. hR, hours of regeneration; dR, days of regeneration. Scales, 300 µm (VC1 images), 500 µm (*gpas* images and A), 150 µm (TMUS and AC-TUB images).

**Table S1.** (A). Gene ontology enrichment for biological processes (BPs) of the 781 up-regulated transcripts in the comparison 7dS vs 1dS (q-val < 0.1). Redundant GO terms were removed from the list by using REVIGO (small similarity allowed). (B) Gene ontology enrichment for biological processes (BPs) of the 814 down-regulated transcripts in the comparison 7dS vs 1dS (q-val < 0.1). Redundant GO terms were removed from the list by using REVIGO (small similarity allowed). (C) Gene ontology enrichment for biological processes (BPs) of the 491 down-regulated transcripts in the comparison 30dS vs 1dS (q-val < 0.1). Redundant GO terms were removed from the list by using REVIGO (small similarity allowed). (D) Upregulated genes at 7dS vs 1dS and at 30dS vs 1dS which belong to known signalling pathways involved in the regulation of stem cell function under CR or fasting. Upregulated transcripts were annotated with KO (KEGG Orthology) and mapped to KEGGs pathways by using KAAS (KEGG Automatic Annotation Server (Moriya et al., 2007). Transcripts and KOs assigned are indicated for each pathway. (E) 41 down-regulated transcripts at 7dS vs 1dS presenting GO term for "protein folding" (GO:0006457). Transcript IDs correspond to IDs from Dresden transcriptome (Planmine) (Rozanski et al., 2019)). Gene symbols and description were assigned by selecting one of the top3 blast hits given by Planmine. Localization and functional type of chaperones is indicated according to Brehme et al. 2014. (F) 21 down-regulated transcripts at 30dS vs 1dS presenting GO term for "protein folding" (GO:0006457). Transcript IDs correspond to IDs from Dresden transcriptome (Planmine; Rozanski et al., 2018). Gene symbols and description were assigned by selecting one of the top3 blast hits given by Planmine. Localization and functional type of chaperones is indicated (Brehme et al., 2014).

**Table S2.** (A) Cluster assignment for the 2070 differential expressed genes (DEGs) found in the X1 at 1dS, 7dS and 30dS. Transcript IDs correspond to IDs from Dresden transcriptome version 4 (Planmine) (Rozanski et al., 2019). Normalized TPMs for each sample, q-values for every pairwise comparison and clustering assignment for all DEGs in X1 are shown. (B) Gene ontology enrichment for biological processes (BPs) of the transcripts from every cluster. Redundant GO terms were removed from the list by using REVIGO (small similarity allowed). No biological processes were found enriched for the clusters 3 and 6. (C) Transcripts from cluster 1 presenting GO term for "protein folding" (GO:0006457). Transcript IDs correspond to IDs from Dresden transcriptome (Planmine). Gene symbols and description were assigned by selecting one of the top3 blast hits given by Planmine. Localization and functional type of chaperones is indicated (Brehme et al., 2014). (D) Differential expression analysis of X1 vs Xins at 1dS, 7dS, 30dS for genes related to proteostasis. Transcript IDs correspond to IDs from Dresden transcriptome (Planmine). Beta values (b) is a biased estimator of the fold change. Positive beta values indicate up-regulation in the X1. q-values represent the false discovery rate adjusted p-value, using Benjamini-Hochberg.

**Table S3.** (A) Gene ontology enrichment for biological processes (BPs) of the 982 up-regulated transcripts in the comparison *cct3A* RNAi vs *gfp* RNAi at 72hR (q-val < 0.05). Redundant GO terms were removed from the list by using REVIGO (small similarity allowed). (B) 34 up-regulated transcripts in *cct3A* RNAi at 72hR presenting GO term for "protein folding" (GO:0006457). Transcript IDs correspond to IDs from Dresden transcriptome (Planmine) (Rozanski et al., 2019). Gene symbols and description were assigned by selecting one of the top3 blast hits given by Planmine. Beta values (b) is a biased estimator of the fold change. Positive beta values indicate up-regulation in the X1. q-values represent the false discovery rate adjusted p-value, using Benjamini-Hochberg. Localization, functional type of chaperones and ATP-dependency is indicated according to Brehme et al. 2014. (C) List of transcripts differentially regulated in *cct3A* RNAi at 72hR which present go term for "DNA damage" and/or "DNA repair". Transcript IDs correspond to IDs from Dresden transcriptome (Planmine). Gene symbols and description were assigned by selecting one of the top3 blast hits given by Planmine. Genes related to "DNA damage" and "DNA repair" were retrieved from the transcripts in the planarian Dresden transcriptome (Planmine) presenting the terms GO:0006974 and GO:0006281 respectively. (D) List of transcripts differentially regulated in *cct3*A RNAi at 72hR which present go term for "endocytosis" (GO:0006897). Transcript IDs correspond to IDs from Dresden transcriptome (Planmine). Gene symbols and description were assigned by selecting one of the top3 blast hits given by Planmine. Genes related to "endocytosis" were retrieved from the transcripts in the planarian Dresden transcriptome (Planmine) presenting the terms GO:0006897.

**Table S4.** (A) Gene ontology enrichment for biological processes (BPs) of the 1431 up-regulated transcripts in the comparison *xbp1/atf6* RNAi vs *gfp* RNAi at 72hR (q-val < 0.05). Redundant GO terms were removed from the list by using REVIGO (small similarity allowed). (B) Gene ontology enrichment for biological processes (BPs) of the 1735 down-regulated transcripts in the comparison *xbp1/atf6* RNAi vs *gfp* RNAi at 72hR (q-val < 0.05). Redundant GO terms were removed from the list by using REVIGO (small similarity allowed).

**Table S5.** (A) List of transcripts related to lipid metabolism differentially regulated in *cct3A* RNAi and/or *xbp1/atf6* RNAi at 72hR. Transcript IDs correspond to IDs from Dresden transcriptome (Planmine). Gene symbols and description were assigned by selecting one of the top3 blast hits given by Planmine. Genes related to "fatty acid beta-oxidation" and "sphingolipid biosynthetic process" were retrieved from the transcripts in the planarian Dresden transcriptome (Planmine) presenting the terms GO:0006635 and GO:0030148 respectively. (B) 146 transcripts related to lipid metabolism down-regulated in *xbp1/atf6* RNAi at 72hR. Transcript IDs correspond to IDs from Dresden transcriptome (Planmine). Gene symbols and description were assigned by selecting one of the top3 blast hits given by Planmine.
